# The Requirement of Ubiquitin C-Terminal Hydrolase L1 (UCHL1) in Mouse Ovarian Development and Fertility

**DOI:** 10.1101/2022.02.08.479454

**Authors:** Morgan F. Woodman, Meghan C.H. Ozcan, Megan A. Gura, Payton De La Cruz, Alexis K. Gadson, Kathryn J. Grive

**Author notes:** Corresponding author*: EMAIL; Kilguss Research Institute, 200 Chestnut St., Room 108, Providence, RI 02903, USA.

## Abstract

Ubiquitin C-Terminal Hydrolase L1 (UCHL1) is a de-ubiquitinating enzyme enriched in neuronal and gonadal tissues known to regulate the cellular stores of mono-ubiquitin and protein turnover. While its function in maintaining proper motor neuron function is well-established, investigation into its role in the health and function of reproductive processes is only just beginning to be studied. Single-cell-sequencing analysis of all ovarian cells from the murine perinatal period revealed that *Uchl1* is very highly expressed in the developing oocyte population, an observation which was corroborated by high levels of oocyte-enriched UCHL1 protein expression in oocytes of all stages throughout the mouse reproductive lifespan. To better understand the role UCHL1 may be playing in oocytes, we utilized a UCHL1-deficient mouse line, finding reduced number of litters, reduced litter sizes, altered folliculogenesis, morphologically abnormal oocytes, disrupted estrous cyclicity and apparent endocrine dysfunction in these animals compared to their wild-type and heterozygous littermates. These data reveal a novel role of UCHL1 in female fertility as well as overall ovarian function, and suggest a potentially essential role for the ubiquitin proteasome pathway in mediating reproductive health.

**Summary sentence:** Ubiquitin C-Terminal Hydrolase L1 (UCHL1) is required for proper ovarian folliculogenesis, estrous cyclicity, and fertility in the female mouse.

## INTRODUCTION

In all mammals, the establishment of the ovarian reserve occurs during fetal or neonatal life, although the timing of the process underlying the formation and the exhaustion of the reserve is species-specific. The oocytes present in the adult human gonad derive from primordial germ cells (PGCs), which migrate into the gonadal ridges and proliferate as oogonia within germ cell cysts during the fifth week of embryonic life [1–4]. This process, known as oogenesis, continues with the oogonia mitotically dividing into primary oocytes, which then initiate meiotic prophase. Following this, oocytes separate by a process called “cyst break down” and become enclosed in primordial follicles consisting of one oocyte arrested in prophase I of meiosis surrounded by a layer of somatic pre-granulosa cells. Upon cyst break down, only one-third of the total oocyte population will survive, while the rest succumb to apoptotic cell death. The remaining oocytes form the entire primordial follicle pool available during a female’s reproductive lifetime and determine her reproductive capacity. This finite population of follicles remain quiescent until follicle activation, at which point they are recruited to undergo maturation and ovulation, or are lost due to follicular atresia [5,6]. Mice experience a highly analogous process of ovarian folliculogenesis, with ovarian reserve development occurring perinatally and follicle maturation utilizing similar mechanisms to those identified in the human ovary [4]. As the ovarian reserve represents the entire procreative potential of a female, it is essential that this population achieve healthy quiescence after birth, allowing for subsets of these oocytes to be activated, recruited, matured, and potentially ovulated after reproductive maturity. To accomplish these goals, not only must these immature oocytes be available in sufficient numbers, but they must also be competent to communicate with their surrounding somatic cells. Both oocyte and somatic compartments must also be appropriately responsive to the effects of steroid hormones and gonadotropins including estrogen, progesterone, follicle stimulating hormone (FSH), and luteinizing hormone (LH) [7], as well as growth factors and signaling molecules [5,8,9].

Proper establishment and function of reproductive processes are dependent on an intricate, yet not fully understood, cascade of events. It has been suggested that the ubiquitin–proteasome pathway plays an important role in many of these processes, including gametogenesis, fertilization, implantation, and embryonic development [10–12]. The ubiquitin-proteasome pathway is responsible for substrate-specific protein degradation and turnover and also plays an important role in eukaryotic cell cycle. Deubiquitinating enzymes (DUBs) catalyze the removal of ubiquitin, regulate ubiquitin-mediated pathways, and are known to contribute to cell growth, differentiation, oncogenesis, and the regulation of chromosome structure [11,13]. Indeed, mutant mice lacking certain ubiquitin protein ligases or ubiquitin-conjugating enzymes lead to defects in meiosis, post-meiotic germ-cell development, and male infertility [14–17]. This omnipresent ubiquitin–proteasome system has evident gonadal implications, but its role in female reproductive processes, specifically ovarian folliculogenesis and endocrine function, remains largely unknown.

We identified one such DUB, Ubiquitin C-Terminal Hydrolase L1 (UCHL1, also known as Protein Gene Product 9.5, PGP9.5) as highly enriched in murine oocytes. Using single-cell-sequencing, our study is one of the first of its kind to be performed on all ovarian cells during the initial establishment of the mouse ovarian reserve. UCHL1 is a multifunctional protein component of the ubiquitin proteasome pathway and was first identified as a member of the ubiquitin carboxyl-terminal hydrolase (UCH) family of DUBs [18,19]. Since then, UCHL1 has been well-documented as an abundant neuronal protein, constituting 5% of the brain-soluble proteins, although it has also been detected in kidneys, breast epithelium, and reproductive tissues [19–22]. Dysfunction of UCHL1 results in impaired motor function and is implicated in variants of common neurodegenerative diseases, such as Parkinson’s and Alzheimer’s diseases [23,24].

While UCHL1 has been primarily studied in the context of neurogenesis, exploration of its role in the ovary is limited. Previous studies using the original Uchl1 *^gad^/_gad_* mutant mouse line [25] have demonstrated that truncation of UCHL1 results in an impaired block to polyspermy, with resulting effects on number of litters and number of pups produced by homozygous animals [26]. This role in the block to polyspermy was also observed in higher mammals. Bovine oocytes which had been subjected to chemical inhibition of UCHL1 followed by *in vitro* fertilization resulted in zygotes with much higher rates of polyspermy compared to controls [11]. Additional work on UCHL1 function in mammalian oocytes similarly utilized chemical inhibitors to test the developmental effects on mouse and rhesus oocytes [27], observing similar consequences on aberrant fertilization, effects which have also been observed in porcine oocytes [12]. Finally, initial studies by Yan-Qiong et al [28] have shown increased UCHL1 expression in morphologically abnormal oocytes in pre-pubertal mouse ovaries, though a mechanism of UCHL1-mediated oocyte survival or loss has not been demonstrated.

Given its highly-enriched expression in mouse oocytes from the time of birth through adulthood, we hypothesized that UCHL1 plays a critical role in ovarian function and female fertility. Our study aimed to better understand the oocyte- and ovary-intrinsic roles of UCHL1 expression, as well as the systemic effects of UCHL1 loss on murine female fertility. In this work, we have demonstrated for the first time that UCHL1 is highly-enriched in all post-natal murine oocytes, and that its loss results in impaired ovarian function, estrous cyclicity, response to hormonal stimulation, and fecundity. Future studies will be critical in dissecting the specific molecular mechanisms of UCHL1, as well as a better understanding of its interacting partners and role in protein homeostasis.

## METHODS

### Animals

C57bl/6 (wild-type) and B6.C-*Uchl1*^gad-J^/J [29] were obtained from the Jackson Laboratory (strain # 024355). B6.C-Uchl1 J/J (“wild-type”), B6.C-Uchl1^gad-J^/J (“heterozygous”), or B6.C-Uchl1^gad-J^/_gad-J_ (“knockout”) mice were generated by mating UCHL1 heterozygous male and female mice. Offspring were genotyped by PCR analysis of ear-snip genomic DNA amplifying the region targeted by homologous recombination using genotyping protocols established by the Jackson Laboratory. All animal protocols were reviewed and approved by Brown University Institutional Animal Care and Use Committee and were performed in accordance with the National Institutes of Health Guide for the Care and Use of Laboratory Animals (# 19-07-0001). All animal protocols were all reviewed and acknowledged by the Lifespan University Institutional Animal Care and Use Committee (# 1668974-1).

### Fertility Trials

To assess female fertility in all genotypes of B6.C-*Uchl1*^gad-J^/J mice, controlled fertility trials were performed. B6.C-Uchl1 J/J, B6.C-Uchl1^gad-J^/J, or B6.C-Uchl1^gad-J^/_gad-J_ females were housed with wild-type C57BL/6J males, both at 6 weeks of age (n=3 for both wild-type and knockout, n=4 for heterozygotes). All animals were housed in single breeding pairs with love huts, and monitored for signs of copulation, pregnancy, and birth of pups. Number and sizes of litters were recorded. As *Uchl1*-null mice develop severe motor dysfunction by 6-7 months of age, all breedings were discontinued at 20 weeks of age, and compared between genotypes.

### Vaginal Cytology and Estrous Cycle Monitoring

Daily vaginal smear cytology was performed over the course of two weeks to determine estrous cyclicity, beginning at 6 weeks of age on virgin females (n=4 for wildtype, n=6 for heterozygotes, and n=5 for knockouts). Briefly, the vagina of each mouse was flushed with a small volume of sterile saline solution and then mixed on a glass slide with Toluidine Blue O dye [30]. Slides were visualized on an EVOS M5000 Fluorescence Imaging System and were classified as in proestrus, estrus, or metestrus/diestrus based on the majority cell type present in the vaginal sample [30]. The percentage of time spent in each sub-stage was quantified per mouse and averaged per genotype. Following estrous cycling, mice were allowed to age to 5 months, at which point mice were sacrificed in proestrus, whole blood was collected for serum analysis, and ovaries collected for analysis of ovary weights versus body weights (n=3 wild-type, 5 heterozygous, 5 knockout animals).

### Ovarian Histology

Ovaries were removed at 1 month of age (n=8 wild-type, 6 heterozygous, 6 knockout animals), or 5 months of age (n=4 wild-type, 2 heterozygous, 4 knockout animals) cleaned of excess fat and bursal sac, and fixed in 1:10 formalin solution overnight. Standard paraffin embedding, 5 μM serial sectioning, and hematoxylin-and-eosin staining were performed [31]. Slides were visualized on an EVOS M5000 Fluorescence Imaging System and images of all fields of a single section were captured. Ovarian follicle counts were quantified using two sections on every fourth slide of sectioned ovary to capture all follicle stages without over-counting of larger follicles (given 8 sections per slide, at 5um per section, every 4^th^ slide represents 160um of distance). Follicle counts were normalized to section area to account for size differences between different ovaries. Follicles were classified by standard protocols [32]. In brief, primordial follicles were defined by one layer of flattened granulosa cells surrounding the oocyte; primary follicles were defined by a single later of cuboidal granulosa cells surrounding the oocyte; secondary follicles were defined by two or more layers of cuboidal granulosa cells surrounding the oocyte; pre-antral follicles were classified as follicles in which antral space had begun to form among the granulosa cells; and antral follicles were classified as follicles in which a complete semi-circular antral space had formed and cumulus-surrounded oocytes were observable.

### Hormonal stimulation and super-ovulation

For hormonal stimulation experiments followed by histology, 1 month old mice were injected intraperitoneally with 5 IU Pregnant Mare Goat Serum (PMSG; Prospec Bio) (n=2 wild-type and 2=knockout animals). Forty-eight hours later, one mouse from each genotype was intraperitoneally injected with 5 IU Human Chorionic Gonadotropin (HCG; Prospec Bio), while the other two mice (one of each genotype) were euthanized and ovaries collected for histology. Twelve hours later, HCG-injected mice were euthanized and ovaries collected for histology. For super-ovulation experiments, mice at 4 weeks of age were super-ovulated by standard protocols [33] (n=13 wild-type, 14 heterozygous, 3 knockout animals). Briefly, mice were injected intraperitoneally with 5 IU Pregnant Mare Goat Serum. Forty-eight hours later, the same mice were intraperitoneally injected with 5 IU Human Chorionic Gonadotropin. Twelve hours later, mice were euthanized and ovaries and oviducts collected. Ovulated cumulus-oocyte complexes were released from the ampulla of the oviduct and counted per mouse. Number of oocytes per mouse were averaged between genotypes and compared.

### Serum hormone analysis

For AMH analysis, whole blood was collected from mice at 80 days of age by cardiac puncture (n=3 wild-type, 7 heterozygous, 6 knockout animals). Blood was collected into serum separator tubes, allowed to clot for 30 minutes, then spun at 3000g for 15 min at 4 °C. The supernatant was collected as serum for analysis. Serum was sent to the University of Virginia Ligand Assay & Analysis Core of the Center for Research in Reproduction for analysis of Anti-Mullerian Hormone serum concentrations. For LH and FSH analysis, blood was collected from 5 month old animals (n=2 per genotype) and serum purified as described. Serum was analyzed as described at the University of Virginia.

### Generation of Ovarian Single Cell Libraries

C57bl/6 ovaries were collected from mice at embryonic day 18.5 (E18.5), and postnatal days (PND) 1, 3, and 5 and pooled to generate pools of 1 million cells per time-point (E18.5 n=47 ovaries, PND1 n=29 ovaries, PND3 n=25 ovaries, PND5 n=8 ovaries). Ovaries were dissociated by incubation in 750 μL 0.25% Trypsin-EDTA for 30 min at 37 °C with rotation. Cells were then spun at 500 g for 5 min, and resuspended in 150 μL 7 mg/mL DNase I and gently pipetted to disperse cells. Cells were incubated for 5 min at room temperature, and then DNase I activity was quenched by addition of 1 mL PBS + 10% Knockout Serum Replacement. Cells were filtered through a 40 μM filter and spun again at 500 g for 5 min. The resulting cell pellets were counted and resuspended in freezing media (70% DMEM, 20% Knockout Serum Replacement, 10% DMSO) and stored at −80 °C in cryotubes.

Cell preparations were then submitted to Genewiz for 10X Genomics scSEQ on the 10X Genomics Chromium System. Library preparation was performed by Genewiz, per the manufacturer’s instructions and as previously described [34]. Libraries were sequenced to an average depth of 394M reads (range 372M-423M); on average, 93% of reads (range 88%-95%) mapped to the reference genome (/gwngsfs/gwngs/data/ref/cellranger/refdata-cellranger-mm10-3.0.0). Initial quality control (Supplementary Figures 2-6) and bioinformatics analyses were conducted at GENEWIZ, LLC. Loupe Cell Browser 2.0.0 was used for downstream analysis of ovarian somatic cells and oocytes at all time points and visualization with t-distributed stochastic neighbor embedding (tSNE) plots.

Data availability: The single-cell RNAseq data have been deposited in the NCBI Gene Expression Omnibus (GEO) and are accessible through accession number GSE186843.

### Single-cell RNA seq (scSEQ) data analysis [35]

The PND0 sample SRX6451401 from GEO Series GSE134339 was downloaded from NCBI SRA onto Brown University’s high-performance computing cluster at the Center for Computation and Visualization. In this published dataset, isolated PND0 ovaries from C57BL/6J mice were cut into pieces and transferred into 0.25 % trypsin/EDTA and collagenase for 6-8 minutes at 37 °C. Following the termination of digestion, the single cell suspension was filtered with a 40 μm mesh strainer and washed twice with PBS containing 0.04 % BSA. Single-cell RNA-seq libraries were prepared using Single Cell 3’ Library and Gel Bead Kit V2 (10x Genomics Inc., 120237, Pleasanton, CA, USA) following the manufacturer’s instructions. This cell preparation is highly analogous to the cell preparation for our single-cell sequencing presented in this study, allowing reasonable analysis of the relationship between *Uchl1* expression and that of other oocyte-specific genes, and for those findings to be applicable to our own. The downloaded fastq files were aligned using Cell Ranger (v 5.0.0, 10x Genomics Inc) count to the mm10 genome. The resulting “filtered feature bc matrix” was used as input for Seurat (v 3.9.9, a software package designed for quality control and analysis of scSEQ data). in RStudio (R v 4.0.2). Seurat [35] was used to select for *Ddx4-positive* (Ddx4 > 0), high-quality (nFeature_RNA > 100, nFeature_RNA < 7000, nCount < 40000, percent mitochondrial genes < 15%) oocytes. To obtain gene counts, the Seurat function “GetAssayData” under the slot “counts” was used. These gene expression counts for *Uchl1* compared to *Sohlh1, Figla*, and *Lhx8* were plotted in Graphpad Prism and a linear regression applied (Supplementary Figure 9).

### Immunofluorescence

Wild-type ovaries were collected at E18.5, PND0, PND1, PND3, 3 weeks, 1 month, 5 months, and 8 months of age (n=2 per time point), cleaned of excess fat, and fixed in 1:10 formalin solution overnight before dehydration and embedding in paraffin by the Brown University Molecular Pathology Core. Fixed ovaries were sectioned at 5 μm onto glass slides by the Brown University Molecular Pathology Core. To stain, sections were de-paraffinized by standard protocols [36] with minor modifications: 3x, 5 minute washes in Safeclear followed by rehydration in 100% ethanol (2x, 5 minutes), 95% ethanol (2x, 5 minutes), 70% ethanol (1x, 5 minutes), water (1x, 5 minutes). Staining was carried out by standard protocols [37]: sections were then incubated in boiling antigen retrieval buffer (10 mM sodium citrate, 0.05% Tween-20, pH 6.0) for 20 minutes and left to cool. Sections were washed 3x, 5 minutes in 1X PBS + 0.1% Triton-X (PBST). Tissue sections were then incubated in blocking buffer [3% Goat Serum (Sigma), 1% Bovine Serum Albumin (Sigma), and 0.5% Triton-X (Fisher Scientific) in 1X PBS] and stained by incubation with primary antibodies against UCHL1 (Proteintech #14730-1-AP, 1:100) and TRA98 (Abcam #ab82527; 1:100) overnight at 4 °C. The following day, slides were washed 3x, 5 minutes in PBST and then incubated with secondary antibodies raised in goat against rat (594 nm) and rabbit (488 nm) at 1:500 for 1 hour at 37 °C. A secondary antibody-only control was included to assess background staining (Supplementary Figure 10B). Sections were further stained with DAPI to visualize nuclei, mounted and analyzed on a Nikon TE2000e inverted epifluorescence microscope. For a given set of antibodies, images were exposed equivalently for all samples from different time points to generate images for relative comparison of intensity over time.

### Timeline of experimental procedures and cohorts

- Single-cell sequencing and validation cohort: C57bl/6 animals were maintained as breeders to produce pups for collection at embryonic day 18.5, and post-natal days 1, 3, and 5. Pups were collected at respective time points for ovary collection and single-cell sequencing. Additional pups from these litters were collected at those same time points, or allowed to age to 3 weeks, 1 month, 5 months and 8 months for immunofluorescence validation of scSEQ results.
- Breeding cohort for fertility trials: heterozygous crosses were maintained to produce pups of all genotypes. Beginning at 6 weeks of age, wild-type, heterozygous, or knockout females were housed with age-matched wild-type males. Animals were allowed to breed until 20 weeks of age at which point they were sacrificed.
- Estrous cycling, folliculogenesis, and serum hormone analysis cohorts: heterozygous crosses were maintained to produce pups of all genotypes. Beginning at 6 weeks of age, wild-type, heterozygous, or knockout females were monitored daily for two weeks by vaginal cytology. These same mice were then retained until the age of 5 months, when they were collected in the proestrus phase of the cycle. Serum was collected for analysis of serum FSH and LH, and ovaries were collected for weights and histological analysis of follicle development. Additional animals which had not been profiled for estrous cyclicity were sacrificed at 1 month for early folliculogenesis analysis or at 2.5 months of age for analysis of serum AMH.
- Hormonal stimulation cohort: heterozygous crosses were maintained to produce pups of all genotypes. Wild-type, heterozygous, or knockout females at 1 month of age were hormonally stimulated with PMSG alone or PMSG followed by HCG, then sacrificed for ovary collection and histological analysis of stimulated ovaries.
- Super-ovulation cohort: heterozygous crosses were maintained to produce pups of all genotypes. Wild-type, heterozygous, or knockout females at 4 weeks of age were hormonally stimulated with PMSG followed by HCG, after which animals were euthanized and cumulus-oocyte complexes collected for analysis of oocyte retrieval outcomes.

### Statistical Analysis

All statistical analyses were performed in GraphPad Prism, and all data assessed for normal distribution. Simple linear regression analyses of single-cell sequencing gene expression were carried out with Pearson correlation co-efficients and p-values calculated. Fecundity analyses were performed using 2-way ANOVA with Šídák’s multiple comparisons tests applied. Follicle count and estrous cyclicity analyses were performed using 2-way ANOVA with Tukey’s multiple comparisons tests applied. Analysis of ovary weights were performed using 1-way ANOVA with Tukey’s multiple comparisons test applied. Serum AMH and LH analyses were also performed using 1-way ANOVA with Tukey’s multiple comparisons test applied.

## RESULTS

### *Uchl1* is highly expressed in oocytes within the mouse ovary, with protein localization in oocytes of all stages

In an effort to identify and profile the dynamic gene expression changes that take place during murine ovarian reserve formation, we collected mouse ovaries at several time points to span the establishment of the ovarian reserve (embryonic day 18.5, E18.5, and postnatal days (PND) 1, 3, and 5). By choosing these time points, we expected to capture oocytes still within ovarian cysts, as well as oocytes undergoing cyst breakdown, and oocytes which are later comprising primordial follicles. For each time point, 4–5,000 cells per mouse were processed through the 10X Genomics Chromium System using standard protocols for single cell RNA sequencing (scSEQ). Libraries were sequenced to an average depth of 1.3 billion reads; on average, 93.1% of reads mapped to the reference genome. After standard data processing, we obtained gene expression profiles for approximately 1100 cells per library (Supplementary Figure 1A, Supplementary Figures 2-6), of which a total of only 70, or about 2%, were germ cells, identified by *Dazl* and *Ddx4* expression (Supplementary Figure 7). While most of the cells in our data set were ovarian somatic cells, this germ cell retention limitation is not unique to this study, with other recent studies also demonstrating comparatively reduced germ cell representation compared to somatic cell representation [38].

We were able to identify globally distinguishing genes in the oocyte population, or genes which are highly and specifically expressed in the germ cells of the developing ovarian reserve. Oocytes selected by expression of the germ cell markers *Dazl* and *Ddx4* (Supplementary Figure 7) were further analyzed to identify novel gene expression dynamics. Our analysis revealed a list of the top 20 oocyte-enriched factors (Supplementary Figure 1A), the vast majority of which are already well-characterized for expression and function, including *Dazl*, meiosis regulators *Sycp1, Sycp3, Syce3, Hormad1*, and *Smc1b* [39]; transcriptional regulators *Figla* [40–42] and *Taf7l* [43]; and retrotransposon regulators *Mov10l1* [44] and *Mael* [45,46]. While these serve as excellent positive controls for the fidelity of our study, our aims to identify novel regulators resulted in further analysis of the 20th most highly-expressed oocyte-expressed gene identified in our study, *Uchl1* (Supplementary Figure 1B). We also observe lower levels of *Uchl1* expression in three specific sub-populations of somatic cells (Supplementary Figure 8), specifically pre-granulosa cells which are *Lgr5*-expression, *Foxl2*-expressing, and *Nr2f2*-expressing [47].

Due to low oocyte yield in our dataset, we also mined existing data sets to independently verify our *Uchl1* oocyte expression data. In the PND0 murine ovarian dataset from Wang et al [48], the authors retained nearly 3000 oocytes for their analysis, allowing us to correlate *Uchl1* expression with other known regulators of oocyte development. Correlation analysis of *Uchl1* with oocyte-specific transcriptional regulators Factor in the Germline Alpha (*Figla*), Spermatogenesis and Oogenesis Specific Basic Helix-loop-helix 1 (*Sohlh1*), and LIM Homeobox Protein 8 (*Lhx8*) revealed significant correlations between the expression of both genes (r=0.7895, r=0.7700, and r=0.7074 respectively; Supplementary Figure 9), further suggesting that *Uchl1* may be playing a key role in oocyte development at the same time as these known master regulators.

UCHL1 has been studied cursorily in the context of oogenesis, though these studies utilized other model systems [12], knockdowns instead of genetic mutants [11,27], or obsolete UCHL1-deficient mouse lines which may not reflect loss of function of the protein [25,26]. Furthermore, UCHL1 has never been studied in the context of ovarian reserve formation, leading us to explore its expression and localization in this context. Wild-type mouse ovaries were obtained from a variety of perinatal (Figure 1) as well as juvenile and adult (Figure 2) time points and immune-stained for UCHL1. Perinatal mouse ovaries were also stained with an antibody against TRA98, an early germ cell marker [49], allowing us to observe the initiation of UCHL1 expression in this population. This staining revealed a protein localization pattern that is consistent with the scSEQ mRNA data, with oocyte-enriched expression through all stages of follicle development. While a low level of *Uchl1* expression was observed in some somatic cells in our scSEQ dataset, this was not observed widely at the protein level, though some protein expression is apparent in the corpora lutea (CL) of adult ovaries (Figure 2, Supplementary Figure 10). From this staining, we also observed UCHL1 co-localization with early oocyte marker TRA98. Interestingly, it is apparent that at E18.5, only a subset of TRA98-positive oocytes are UCHL1-positive, suggesting an intriguing possibility that UCHL1 may represent a part of a survival signature in oocytes, as it is expressed in a subset of oocytes at the peak of perinatal oocyte attrition [3–6], and oocytes which remain after this wave of attrition are nearly all UCHL1-positive (Figure 1). This ubiquitous, oocyte-enriched UCHL1 expression remains throughout juvenile and adult life, including in reproductive-aged 8 month old mice (Figure 2). Importantly, and as expected, UCHL1 knockout ovaries have no detectable UCHL1 expression (Supplemental Figure 10A).

**Figure 1.**
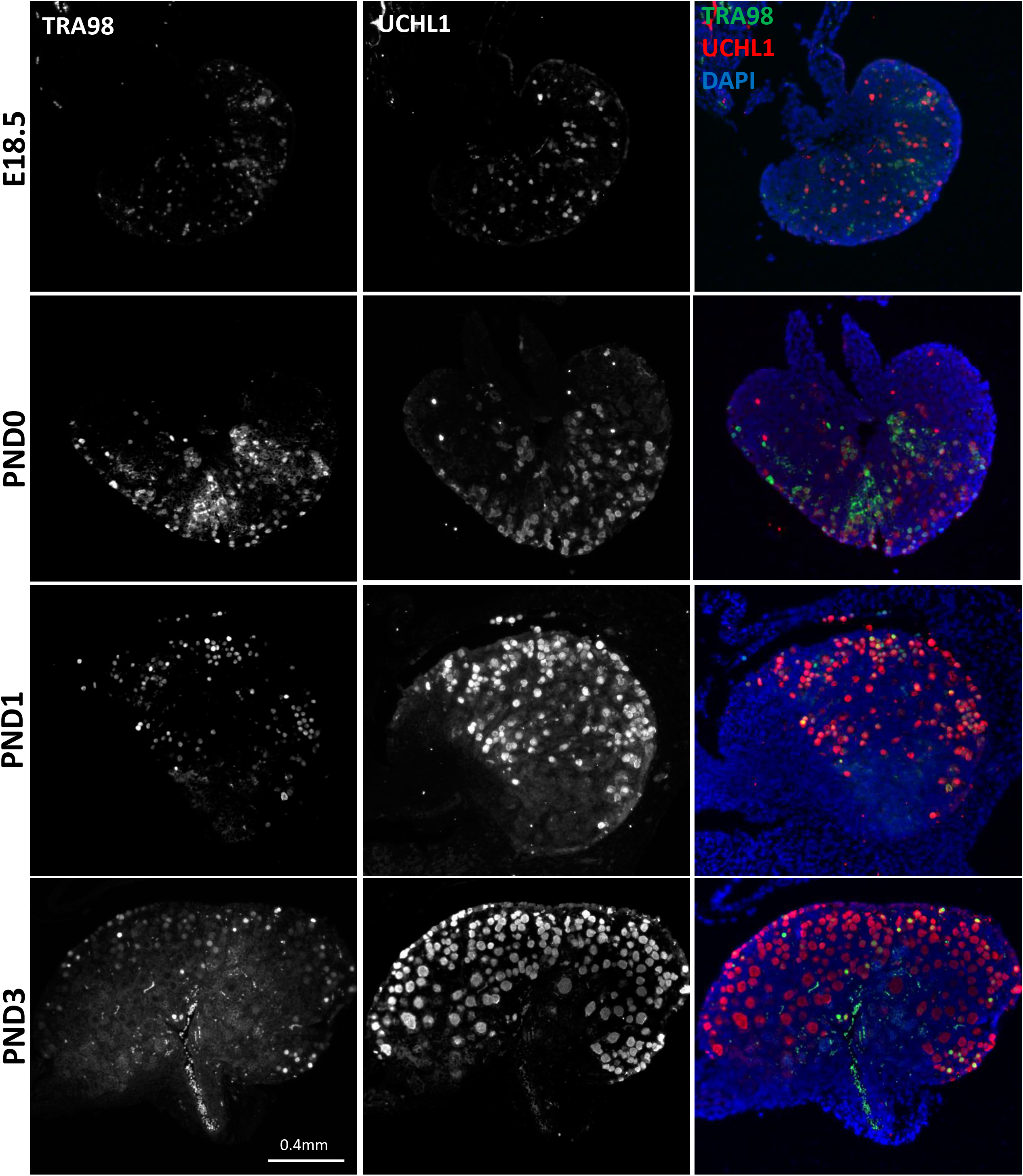
UCHL1 protein expression in perinatal mouse ovaries. Ovaries were collected from mice at E18.5, PND0, PND1, and PND3, paraffin-embedded, sectioned in 5 μm sections, and stained with antibodies against UCHL1 (red) and early oocyte marker TRA98 (green), as well as DAPI (blue).

**Figure 2 -.**
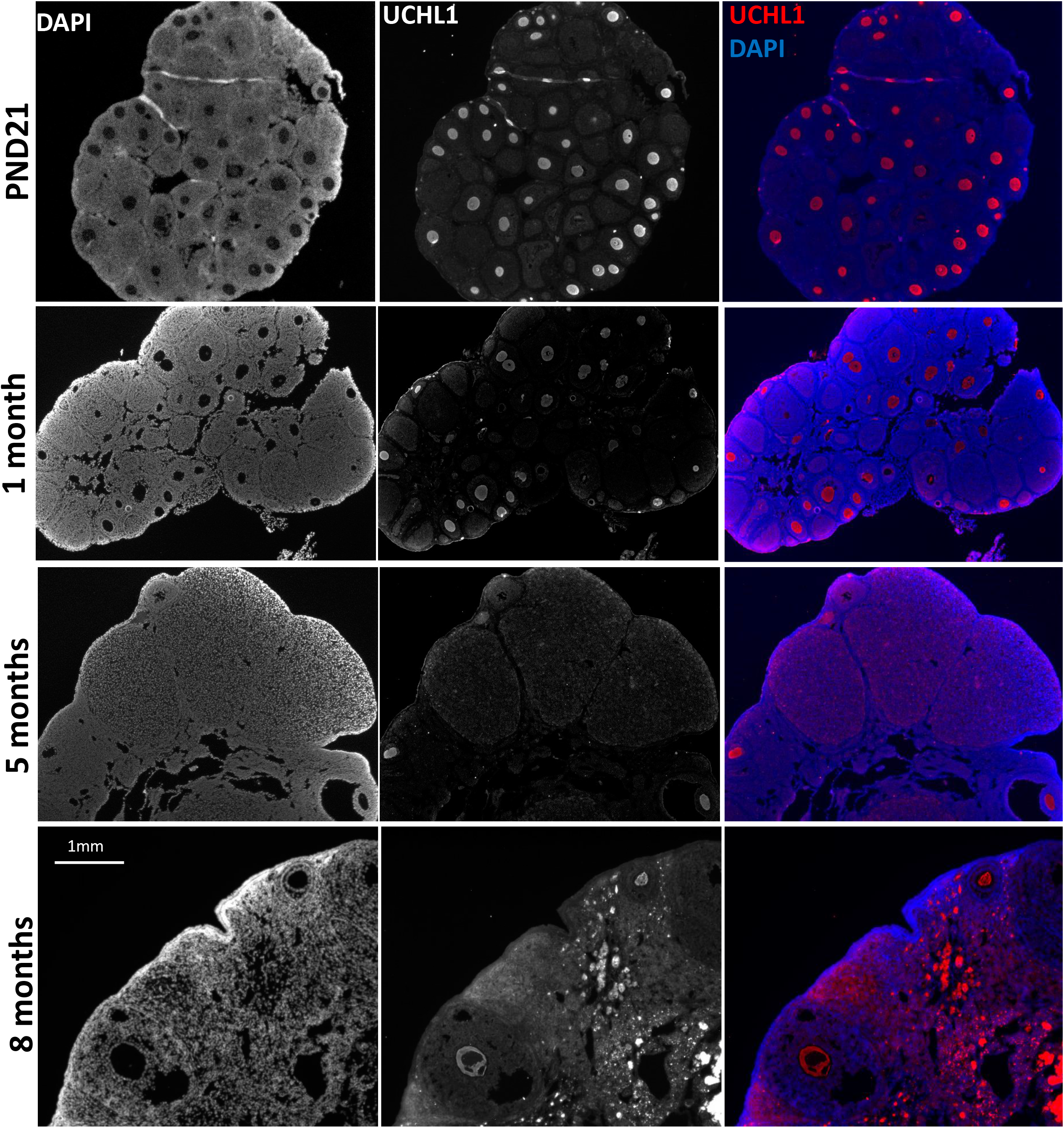
UCHL1 protein expression in juvenile and adult mouse ovaries. Ovaries were collected from mice at PND21, 1 month, 5 months, and 8 months, paraffin-embedded, sectioned in 5 μm sections, and stained with antibody against UCHL1 (red) as well as DAPI (blue).

### UCHL1 knockout mice are severely sub-fertile, with heterozygous mice exhibiting a less severe reduction in fertility

To better understand the role that UCHL1 may be playing in mammalian oocyte development, we obtained a UCHL1-heterozygous mouse line (B6.C-Uchl1^gad-J^/J), which we bred to obtain all genotypes: B6.C-Uchl1 ^J^ /J (“wild-type”), B6.C-Uchl1^gad-J^/J (“heterozygous”), or B6.C-Uchl1^gad-J^/^gad-J^ (“knockout”). Mice of all genotypes were paired with verified C57BL/6J male breeders in single-breeding-pair cages at 6 weeks of age. Mice were monitored for the presence of copulation plugs and normal mating behavior, evidence of pregnancy, the birth of pups, and the number of live pups per litter. Knockout mice appear to breed normally with repeated observation of copulation plugs (data not shown). As UCHL1 knockout mice develop severe motor neuron dysfunction between 6-7 months of age, breedings were only maintained until 5 months of age, at which time all mice were euthanized. Severe sub-fertility in knockout females was demonstrated with significantly fewer litters than wild-type animals (0.667 litters vs. 4.333 litters, respectively, p=0.042). Knockout animals also produce significantly fewer pups per litter than wild-type animals (2.500 pups v.s 8.769 pups, respectively, p=0.0002), as well as fewer pups per litter than heterozygous animals (2.500 pups v.s 6.625 pups, respectively, p=0.008). Heterozygous animals also demonstrate a non-significant trend toward reduced litters and litter sizes (Figure 3).

**Figure 3.**
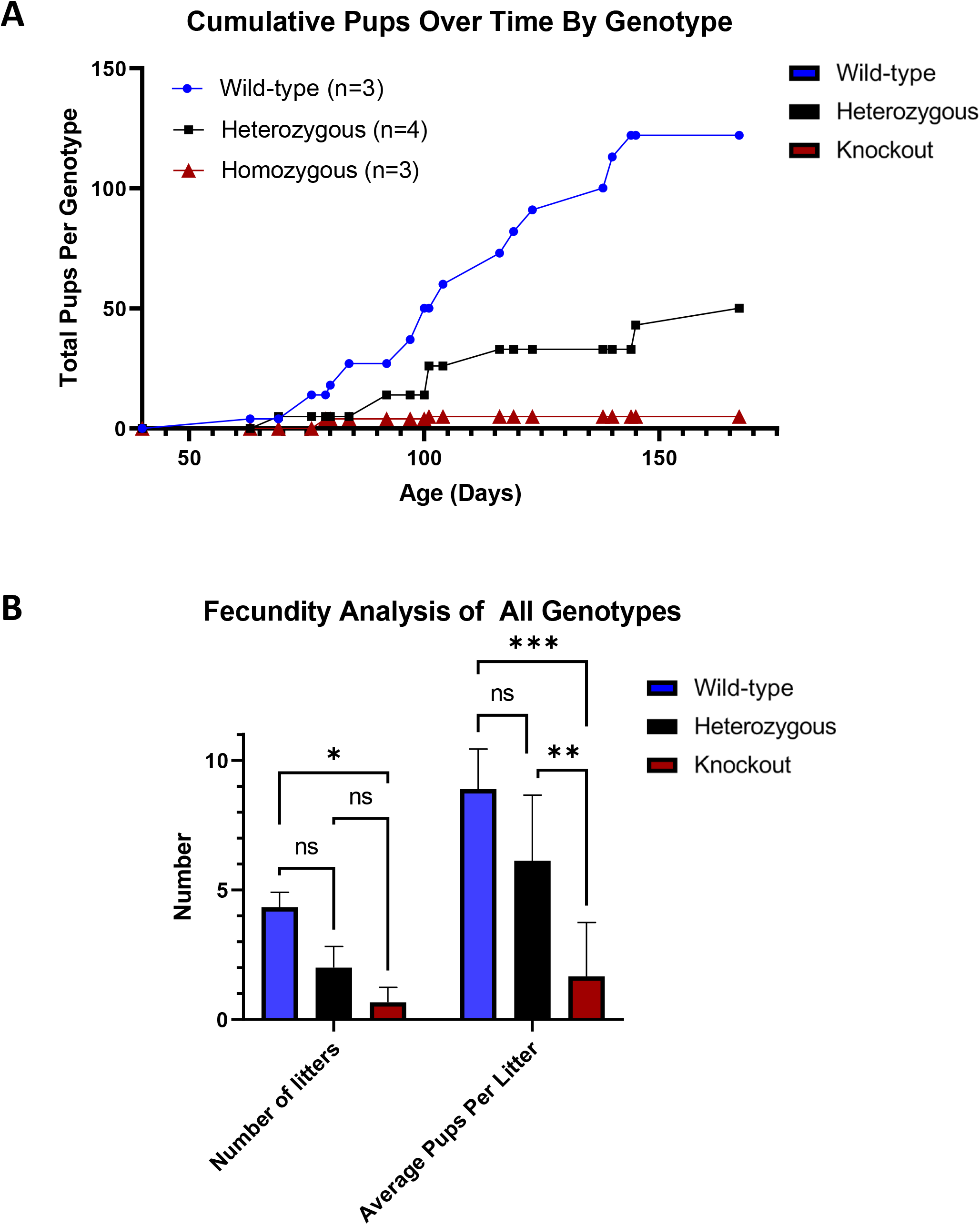
UCHL1 knockout mice exhibit severely reduced fertility with fewer litters produced and fewer pups per litter. A) Cumulative pups resulting from the fertility study were plotted for each genotype. B) Number of litters and pups per litter were graphed for all breeders in 3A.

### UCHL1 knockout mice demonstrate estrous cyclicity disruption as well as reduced response to ovarian stimulation

The severe subfertility seen in knockout animals could be correlated with disrupted estrous cyclicity, reduced follicle abundance or recruitment, and reduced ovulations in these animals. Consistent with this hypothesis, estrous cyclicity analysis in all genotypes of mice revealed significantly disrupted cycles in both knockout and heterozygous animals. Mice were assessed for vaginal cytology daily for two weeks starting at 40 days of age, and percentage of time spent in each cycle stage was averaged between animals in a genotype. Interestingly, both heterozygotes and knockouts spent significantly less time in the follicular phase of proestrus (p=0.0145 and p=0.0085 respectively) and spent significantly more time in the secretory, luteal phase of diestrus (p=0.0434 and p=0.0456 respectively) than wild-type animals (Figure 4A). Additionally, knockout animals trended towards increased time spent in estrus, with several animals spending greater than 5 days in uninterrupted estrus, though this average did not reach statistical significance.

**Figure 4.**
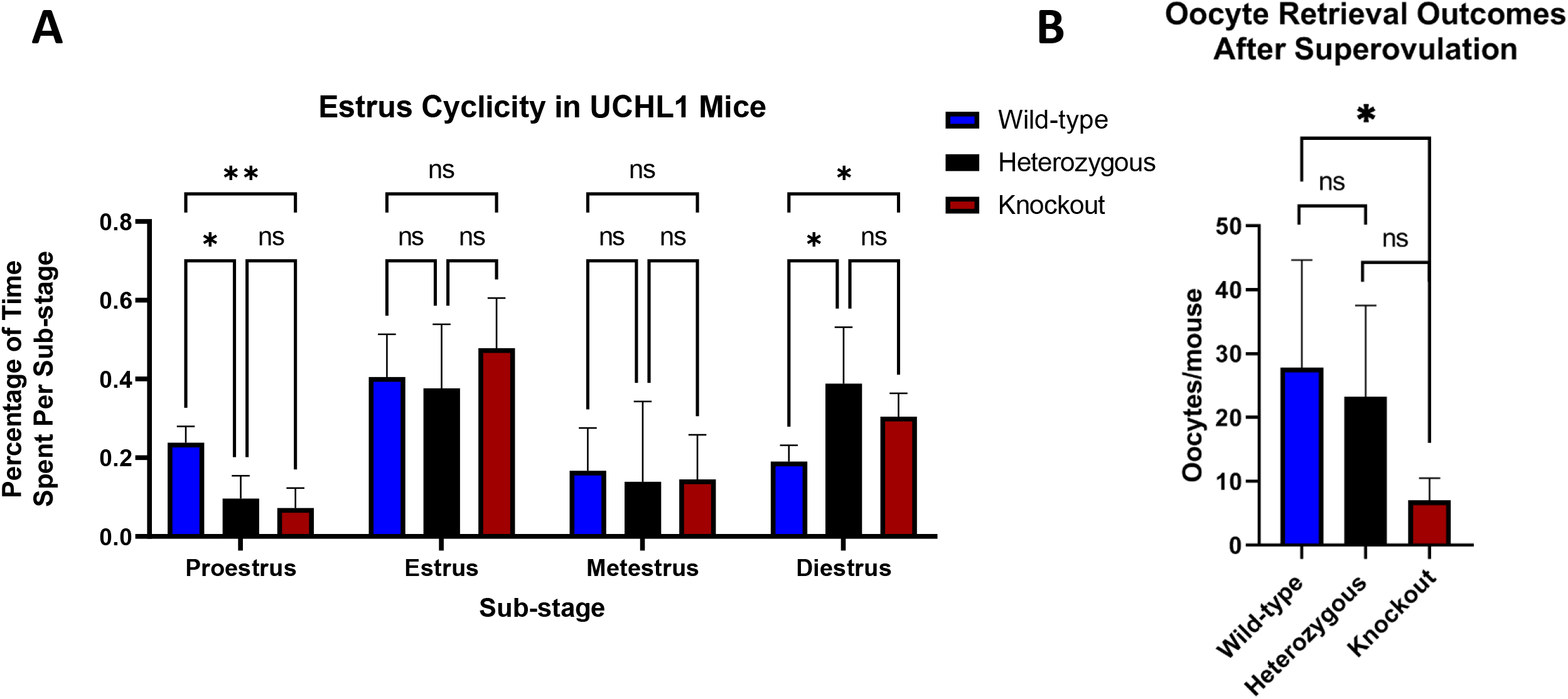
Hormonal cyclicity and response are significantly affected in knockout females. A) Mice were assessed for vaginal cytology daily for two weeks starting at 40 days of age, and percentage of time spent in each cycle stage was averaged between animals in a genotype. B) At 28 days of age, animals of all genotypes were super-ovulated using standard PMSG and HCG protocols. Retrieved oocytes were collected from each animal, counted, and averaged per genotype.

In addition to impaired hormonal cyclicity, knockout animals are also significantly less responsive to ovulation induction (Figure 4B). At 28 days of age, animals of all genotypes were hormonally stimulated using PMSG and HCG, and cumulus-oocyte complexes were collected from the ampullas of all mice. Significantly fewer oocytes were recovered from knockout animals compared to their heterozygous or wild-type counterparts (an average of 7, 23.2, and 27.8 oocytes per animal, respectively, p=0.050), despite ovarian follicle counts appearing indistinguishable at this same time point (Figure 5A). To gain a better understanding of where in the ovulatory process a block to ovulation may be experienced, 1 month old wild-type and knockout animals were stimulated with PMSG alone to assess development of follicles to the antral stage, or with PMSG followed by HCG to assess follicle luteinization. As expected, given that knockout animals are able to ovulate a reduced number of oocytes and even produce small numbers of pups, pre-antral and antral follicles are present following PMSG stimulation, though at potentially slightly delayed stages of maturation compared to wild-type littermates. Interestingly, however, we do also observe the presence of luteinized unruptured follicles (LUFs) [50] in knockout ovaries following both PMSG and HCG stimulation, which are not observed in hormonally stimulated wild-type littermates (Supplementary Figure 11A-D). Interestingly, we also observe significantly elevated LH levels in knockout animals during the follicular phase of proestrus in contrast to wild-type and heterozygous littermates (p=0.0.266 and p=0.0474, respectively, Supplementary Figure 11E), and non-significant elevations in both FSH and the LH:FSH ratio (Supplementary Figure 11F,G) again supporting a phenotype of endocrine dysfunction in these animals.

**Figure 5.**
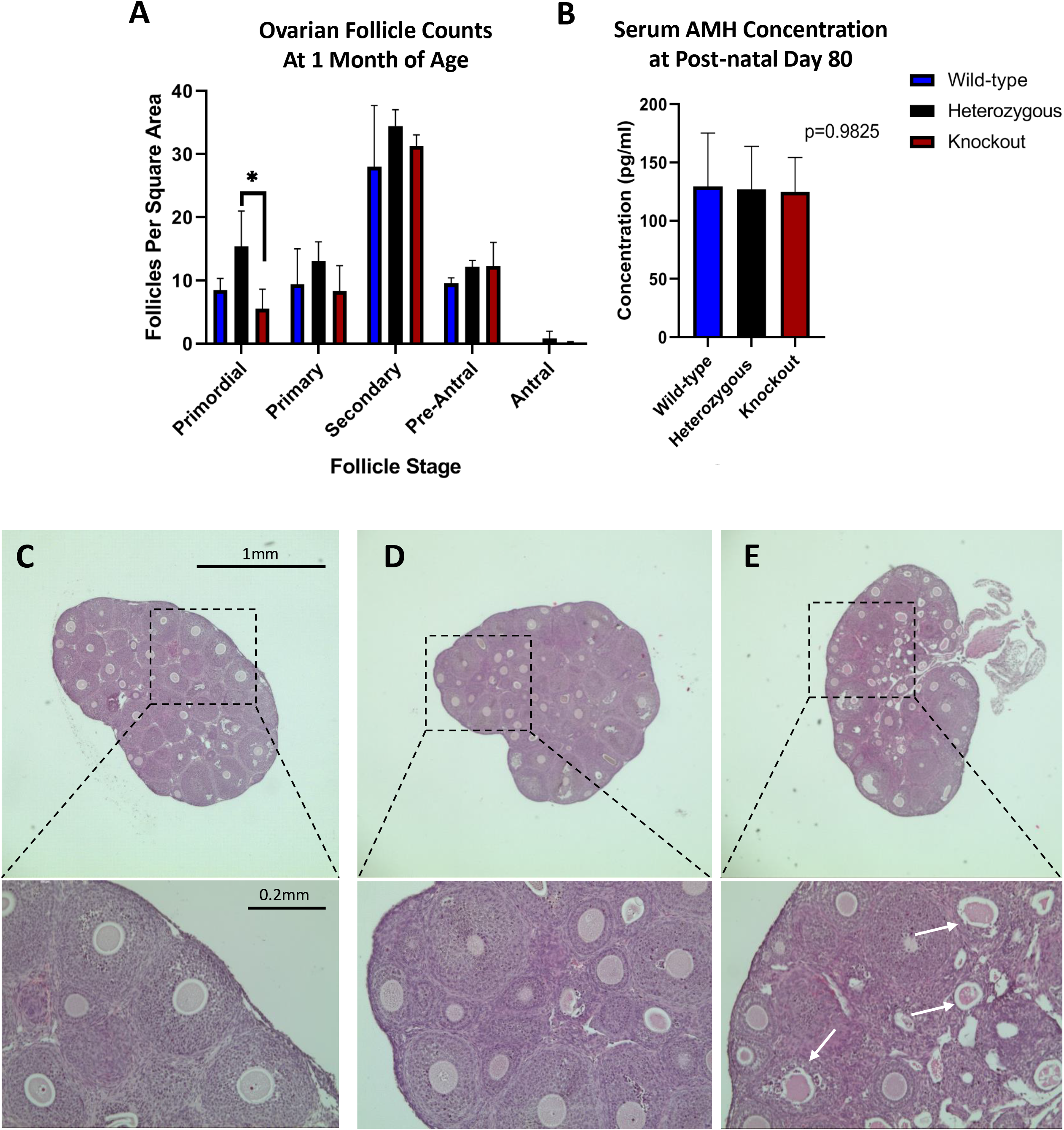
Follicle abundance and AMH levels are initially similar between all genotypes of mice. A) Ovaries from female mice at 1 month of age were collected, paraffin-embedded, and stained by standard hematoxylin and eosin protocols. Ovarian follicle counts were quantified using two sections on every fourth slide of sectioned ovary to capture all follicle stages without over-counting of larger follicles. Follicle counts were normalized to the section area to account for size differences between different ovaries. B) Follicle abundance is consistent with serum AMH, which are not significantly different among the genotypes at 2.5 months of age. C) Representative image of a 1-month-old wild-type ovary with high magnification inset. D) Representative image of a 1-month-old heterozygous ovary with high magnification inset. E) Representative image of a 1-month-old knockout ovary with high magnification inset. White arrows denote degenerating or atretic follicles.

### UCHL1-heterozygous and -knockout ovaries contain normal follicle densities early in life and produce normal Anti-Mullerian Hormone, with ovarian function becoming compromised by middle age

To better understand the ovary-intrinsic effects of UCHL1 loss, ovaries from all genotypes of mice were collected at 1 month and 5 months of age and ovarian follicle counts were quantified. Follicle counts were normalized to the section area to account for size differences between different ovaries. Ovarian weights were also obtained from 5-month-old animals and compared to body weight as a readout of ovarian volume.

At 1 month of age, all genotypes possess similar densities of all follicle stages, suggesting initially normal development and maturation of ovarian follicles, despite qualitative appreciation of increased degenerating or atretic follicles in knockout ovaries (Figure 5A, C-E). While heterozygotes possessed significantly more primordial follicles than knockout animals per unit area at this age (p=0.0308), this effect was not observed in wild-type or knockout animals. As an independent assessment of ovarian follicle endowment, additional mice were collected at 2.5 months, with whole blood collected for serum analysis of Anti-Mullerian Hormone (AMH). AMH is currently the gold standard for measuring oocyte endowment, as it is produced by the granulosa cells of growing follicles [51]. Consistent with our 1 month follicle density data, no differences were observed in AMH serum concentrations between genotypes at 2.5 months of age (Figure 5B).

Interestingly, however, UCHL1 knockout ovaries begin to demonstrate ovarian dysfunction just a few months later, with decreased ovarian volume (Figure 6B,C) as well as altered follicle densities compared to wild-type and knockout animals (Figure 6A, D-F). While primordial follicle densities are the only significantly enriched follicle stage in 5-month-old knockout ovaries (p=0.0206 vs. wild-type animals, and p=0.0443 vs. heterozygous animals), densities of all follicle stages trend towards greater enrichment in knockout ovaries. This is surprising, given the reduced ovarian volume at this time, with knockout ovaries being significantly smaller than wild-type ovaries when normalized for body weight (p=0.050; Figure 6B,C). While this is an unexpected result given the retained follicle density in these animals, these data suggest a potential deficit in oocyte maturation or folliculogenesis.

**Figure 6.**
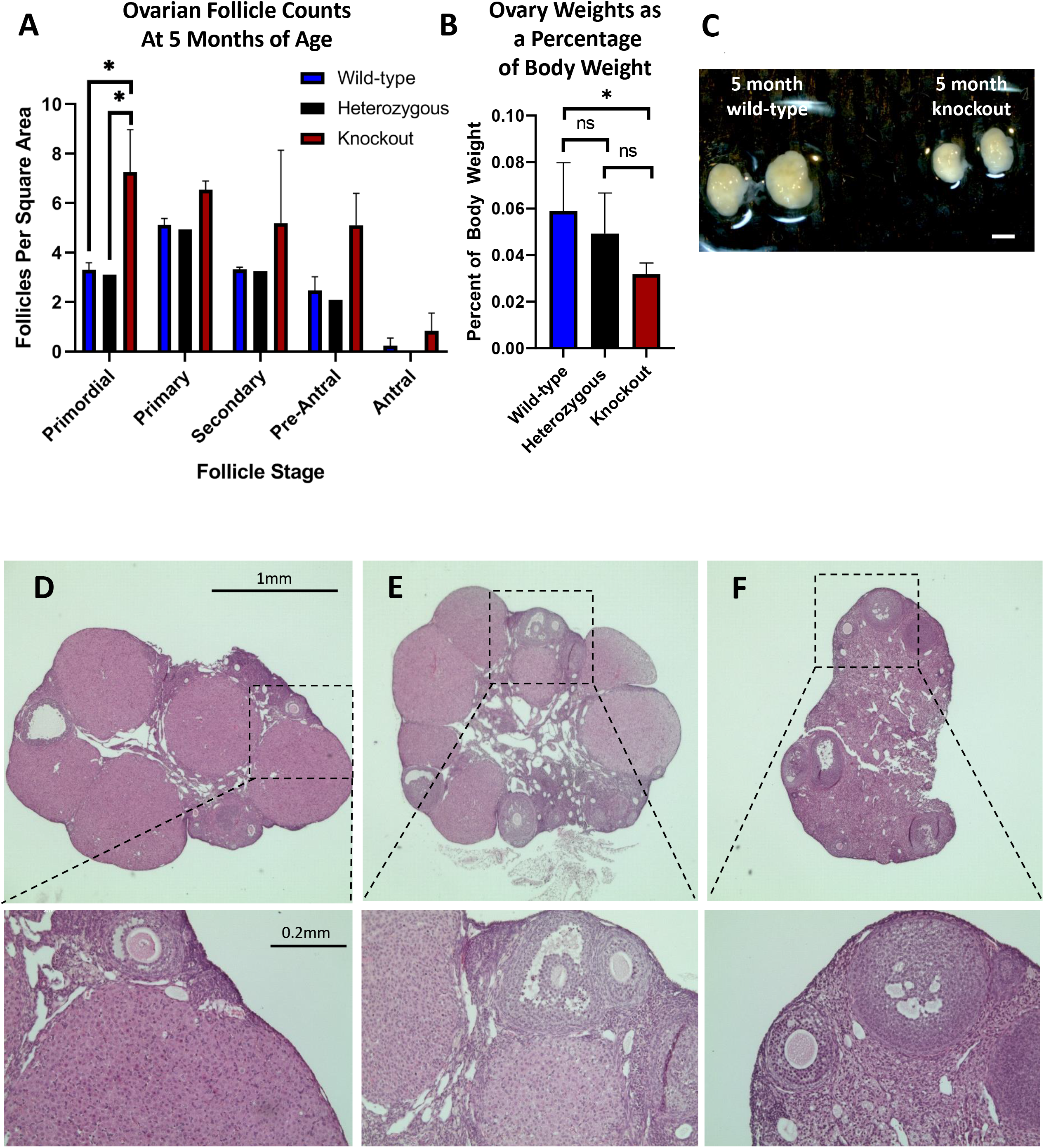
Follicle abundance and ovarian volume is significantly altered in knockout ovaries during mid-reproductive age. A) Ovaries from female mice at 5 months of age were collected, paraffin-embedded, and stained by standard hematoxylin and eosin protocols (n=4 wild-type, 2 heterozygous, 4 knockout animals). Ovarian follicle counts were quantified using two sections on every fourth slide of sectioned ovary to capture all follicle stages without over-counting of larger follicles. Follicle counts were normalized to the section area to account for size differences between different ovaries. B) Ovary weights as a percentage of body weight were averaged at 5 months of age from animals of all genotypes C) Representative image of whole ovaries obtained from 5-month-old wild-type and knockout animals, demonstrating a visible difference in ovarian volume. D) Representative image of a 5-month-old wild-type ovary with high magnification inset. E) Representative image of a 5-month-old heterozygous ovary with high magnification inset. F) Representative image of a 5-month-old knockout ovary with high magnification inset.

## DISCUSSION

The mammalian ovarian reserve directly impacts ovarian function and endocrine balance, making the health of this cell population not only essential for fertility but also female health-span and lifespan. The regulation of oocyte endowment, development, maturation, and resulting endocrine function rely on myriad molecular mechanisms to regulate these developmental transitions, including transcriptional regulation, post-transcriptional and translational mechanisms, and protein homeostasis. As oocytes must remain arrested in the ovary from the time of initial perinatal quiescence to the time of activation weeks to months later in the mouse, or years to decades later in the human, ensuring adequate protein supply, homeostasis, and turnover is likely essential for the developmental success of this population. UCHL1 is a deubiquitinating enzyme known to regulate free cellular mono-ubiquitin, turnover of ubiquitin pro-proteins, and protein abundance and stasis [19]. In this work, we have determined the effects of UCHL1 loss on mouse ovarian development and function, finding an essential role for UCHL1 in the proper execution of mouse folliculogenesis, with loss resulting in severely impaired fertility and hormone responsiveness of the ovary. Interestingly, recent work has also demonstrated a genetic relationship between known oogenesis “master regulators” FIGLA or SOHLH1 and UCHL1 with *Uchl1* mRNA expression reduced nearly 50% in *Figla*-knockout ovaries (p=3.4xe^−13^) and nearly 90% *Sohlh1*-knockout ovaries (p=6.3xe^−133^) [52]. Significant change in *Uchl1* expression was not, however, observed in *Lhx8*-knockout ovaries, despite the expression correlation observed in our analysis. Overall, this speaks to a network of oogenesis-specific regulators of a variety of functions, working in concert to promote establishment and maintenance of a healthy and functional ovarian reserve.

Our finding of reduced litter numbers and litter sizes in B6.C-Uchl1^gad-J^/^gad-J^ ovaries is consistent with previous work using the original Uchl1 *^gad^/_gad_* truncation mutant, but contrasts with other findings from this study. Sekiguchi et al [26] found that while Uchl1 *^gad^/_gad_* animals were significantly sub-fertile, they did not possess visually abnormal ovaries, though follicle abundance was not quantified, and were able to ovulate similar numbers of oocytes compared to wild-type animals when hormonally stimulated. Instead, the authors attribute the observed sub-fertility to a loss of the block to polyspermy, resulting in failed embryo development. Indeed, B6.C-Uchl1^gad-J^/^gad-J^ animals may suffer similar effects on fertilization and early embryo development, though our study also elucidates the ovary-intrinsic effects of UCHL1 loss. The observed differences in phenotype between these two mouse lines may be a result of differences in the site of mutation, as the Uchl1 *^gad^/_gad_* mutation was produced by an in-frame deletion causing UCHL1 truncation of 42 residues [25]. In contrast, the currently available B6.C-Uchl1^gad-J^/^gad-J^ mutation resulted from a spontaneous mutation in a BALB/cJ mouse line at Jackson Laboratory [29]. Back-crossing of this mutation onto a pure C57BL/6J background revealed the mutation to ablate a large intragenic region, ultimately resulting in the truncation of the final 78 residues of the protein. Phenotypic differences may also result from the contribution of different strain backgrounds, as the Uchl1 *^gad^/_gad_* animals studied in the Sekiguchi work were autosomal recessive F1 hybrid mutants resulting from a cross of CBA and RFM animals, while B6.C-Uchl1^gad-J^/^gad-J^ animals are maintained on a pure C57bl/6J background.

Perhaps most intriguing is that UCHL1 knockout ovaries at 1 month of age initially appear to contain comparable follicle densities to their wild-type and heterozygous counterparts, though many of these follicles appear to be morphologically degenerative and likely of poor quality. During mid-reproductive life, despite reduced ovarian size, knockout ovaries have significantly increased follicle densities compared to age-matched ovaries of other genotypes, suggesting a potential delay in follicle maturation or lack of sensitivity to atresia, resulting in follicle retention. Indeed, even when hormonally stimulated, knockout animals at 1 month of age ovulate fewer oocytes than their littermates of other genotypes, suggesting reduced responsiveness to normal hormonal stimuli. This hypothesis gains further support from the observation that knockout animals do not estrous cycle normally, with significantly reduced time spent in the follicular phase, and significantly more time spent in the luteal phase. Finally, analysis of ovaries following PMSG stimulation alone or in combination with HCG demonstrate ovulatory failure in knockout animals, with many follicles developing to the antral stage following PMSG, but oocytes being retained in unruptured luteinized follicles following an HCG trigger. These LUFs are not observed in wild-type ovaries stimulated with the same protocol, suggesting that while some oocytes are able to be ovulated from knockout ovaries, there is a partial anovulatory phenotype in these animals [50], which will be explored in future studies. Interestingly, we observed persistently elevated LH levels in knockout animals during proestrus, in contrast to that seen in their age- and stage-matched wild-type and heterozygous littermates. Furthermore, knockout mice trend toward higher FSH levels, with non-significantly elevated LH:FSH ratios compared to wild-type and heterozygous littermates, which is a phenotype often associated with Polycystic Ovary Syndrome (PCOS) [53,54] or menopause [55]. Indeed, mice which overexpress Luteinizing Hormone Subunit Beta (LHβ) also experience reduced ovulations, extended luteal phase, and reduced fertility [56], much like that which is observed in UCHL1 knockout females, making our observations consistent with endocrine disruption in other models.

These investigations are not without their limitations. We cannot rule out that loss of UCHL1 expression in the brain contributes to these observed endocrine effects, as the hypothalamic-pituitary-ovarian axis creates an inextricable feedback loop between the function of the ovary and the endocrine function of the brain and pituitary gland. Given that UCHL1 is expressed in gonadotropes of the pituitary gland [57], the lack of robust hormone responsiveness in knockout animals is not entirely surprising, and these effects are difficult to parse apart from ovarian effects, as the axis is complex and inter-dependent. Interestingly, other mutant mice with estrous cyclicity defects, including a short proestrus phase, include gonadotropin-releasing hormone (*mGnRH*) mutants [58] as well as DMX Like 2 (*Dmxl2*) mutants, which also affect GnRH signaling [59]. The shorter proestrus phase in these mutant animals, much like that observed in the UCHL1 mutants in this study, suggests that disrupted GnRH production, signaling, and responsiveness could contribute to the altered estrous cyclicity and reduced response to ovulation induction we observe. This is particularly relevant, as GnRH is known to stimulate the pituitary gland to produce follicle stimulating hormone and luteinizing hormone, which are essential for ovulation [60]. Future studies utilizing an oocyte-specific UCHL1-conditional mouse line will assist us in the endeavors to resolve neuroendocrine versus ovarian effects of UCHL1 loss, and will allow us to better assess reproductive fitness in animals which do not suffer premature motor neuron decline and death. UCHL1 loss may also have a more direct effect on oocyte health and developmental potential by directly regulating the protein abundance, composition, and turnover in both resting and maturing oocytes. Loss of UCHL1 in Uchl1 *^gad^/_gad_* oocytes has been shown to reduce free mono-ubiquitin in these cells, likely directly affecting the ability of these oocytes to mature and grow properly [11]. Future studies will further elucidate the underlying biochemical mechanisms of UCHL1 influence in these cells, though the literature provides lessons from the demonstrated role of UCHL1 in the quiescence and differentiation of developing male germ cells. Studies of testis development and function in the original UCHLl1 *^gad^/_gad_* animals revealed that while testis volume is initially normal at 12-weeks-of-age in homozygous mutants, testis volume significantly decreased with age, with an approximately 30% reduction in size by 25 weeks of age, accompanied by reduced spermatogonial stem cell (SSC) proliferation at this same time [61]. This testicular phenotype is much like that we observe in the ovary, with initially normal size, but loss of volume with age. Interestingly, follow-up studies from these authors demonstrated an increase in premeiotic germ cells in juvenile UCHLl1 *^gad^/_gad_* mice, despite reduction in all stages of germ cells later in life, suggesting that UCHL1 may help to regulate both sensitivity to apoptosis as well as developmental competence [62]. If this were to be true for ovarian populations as well, it may explain why we observe degenerating follicles in UCHL1 knockout animals, but without catastrophic loss of follicle abundance. Furthermore, recent studies from Alpaugh et al utilizing a different UCHL1 mouse line, Uchl1^tm1Dgen^, demonstrated that UCHL1-deficient SSCs are less competent to repopulate busulfan-ablated testes, exhibiting only about 20% regenerative capacity compared to 60% in their wild-type counterparts [63]. The authors posit that this lack of developmental potential may result from impaired mitochondrial metabolism of UCHL1-deficient SSCs, and directly result from altered mitochondrial protein turnover. These findings are consistent with recent work utilizing chemical inhibition of UCHL1 in *in vitro* matured mouse oocytes, demonstrating aberrant maturation resulting from altered mitochondrial metabolism [64].

It will be enlightening to explore the intersection of germ cell protein homeostasis, cell metabolism, and developmental competence in future studies. By better understanding the ways in which UCHL1 functions in the development of the oocytes and, therefore, the ovary, we will better understand the ways in which the ubiquitin-proteasome pathway can contribute to protein turnover in highly dynamic cells, as well as govern the ability of those cells to differentiate and develop properly.

## ACKNOWLEDGEMENTS

The authors would like to thank The Program in Women’s Oncology of Women and Infants Hospital and Swim Across America. This work was also supported by University of Virginia Center for Research in Reproduction Ligand Assay and Analysis Core supported by the Eunice Kennedy Shriver NICHD Grant R24 HD102061. We are grateful to the NICHD for their generous support through award 1F31HD097933 to MAG. Part of this research was conducted using computational resources at the Center for Computation and Visualization at Brown University. The authors thank the Freiman and Ribeiro laboratories for their helpful feedback on this project.

## AUTHOR CONTRIBUTIONS

All writing of the manuscript was done by MFW and KJG. Experimental designed was performed by MFW, MCHO, and KJG. Experiments for the manuscript were carried out by MFW, MCHO, PD, AG, and KJG. Analysis for the manuscript was performed by MFW, MAG, PD, and KJG.

## DATA AVAILABILITY

The scSEQ data have been deposited in the NCBI Gene Expression Omnibus (GEO) at https://www.ncbi.nlm.nih.gov/geo/query/acc.cgi?acc=GSE186843 and are accessible through accession number GSE186843.

## CONFLICT OF INTEREST

The authors declare that there are no conflicts of interest.

## FUNDING

This work was funded in part by the NIH P20 GM121298 (COBRE for Reproductive Health) grant as well as funding from Swim Across America.

## FIGURE LEGEND

**Supplementary Figure 1** – *Clustering of single-cell data and Uchl1 expression*. A) tSNE representation of all cells from the four time point libraries of origin with oocytes denoted in inset. The top 20 most highly expressed oocyte-specific genes are shown to the right of the plot. Log2 fold change above somatic cell expression is denoted by the color bar. B) tSNE plot of *Uchl1* expression in all cells.

**Supplementary Figure 2** – *Fragment analysis of single cell sequencing libraries*. Concentrations and fragment electrophoresis of all libraries.

**Supplementary Figure 3** – *Quality control analysis of E18.5 ovarian single sequencing library*. Top) Histogram of all fragments and peaks from E18.5 library. Bottom) Regional histogram demarcating the sequenced fragments between 100-1000bp.

**Supplementary Figure 4** – *Quality control analysis of PND1 ovarian single sequencing library*. Top) Histogram of all fragments and peaks from PND1 library. Bottom) Regional histogram demarcating the sequenced fragments between 100-1000bp.

**Supplementary Figure 5** – *Quality control analysis of PND3 ovarian single sequencing library*. Top) Histogram of all fragments and peaks from PND3 library. Bottom) Regional histogram demarcating the sequenced fragments between 100-1000bp.

**Supplementary Figure 6** – *Quality control analysis of PND5 ovarian single sequencing library*. Top) Histogram of all fragments and peaks from PND5 library. Bottom) Regional histogram demarcating the sequenced fragments between 100-1000bp.

**Supplementary Figure 7** – *tSNE plots of oocyte markers in single-cell data. Dazl* (A) and *Ddx4* (B) were used to identify the oocyte population within the data set. Log2 fold change above somatic cell expression is denoted by the associated color bar.

**Supplementary Figure 8** – *tSNE plots of pre-granulosa cell markers in single-cell data. Lgr5, Foxl2*, and *Nr2f2* were used to identify the somatic cell sub-populations within the data set.

**Supplementary Figure 9** – *Uchl1 expression is highly correlated with expression of other oocyte regulators at PND0*. PND0 oocyte gene expression was mined from the ovarian single-cell data by Wang et al [48]. Normalized gene expression counts for *Uchl1* compared to *Sohlh1, Figla*, and *Lhx8* were plotted in Graphpad Prism and a linear regression applied. Correlation analysis of *Uchl1* with oocyte-specific transcriptional regulators (A) *Figla*, (B) *Sohlh1*, and (C) *Lhx8* revealed significant correlations between the expression of both genes.

**Supplementary Figure 10** – *Specificity controls for immunofluorescence staining*. (A) Ovaries from wild-type and knockout animals at 1 and 5 months of age were paraffin-embedded, sectioned in 5 μm sections, and stained with antibody against UCHL1 (red) as well as DAPI (blue). (B) Ovaries from wild-animals at 5 months of age were paraffin-embedded, sectioned in 5 μm sections, and stained with antibody against UCHL1 (red) as well as DAPI (blue), or stained with secondary antibody only as a negative control.

**Supplementary Figure 11** – *Follicle histology following PMSG or PMSG + HCG ovarian stimulation as well as endogenous gonadotropin levels*. (A) Representative image of wild-type follicles 48 hours after PMSG stimulation. (B) Representative image of knockout follicles 48 hours after PMSG stimulation. (C) Representative image of wild-type corpora lutea following PMSG + HCG stimulation. (D) Representative image of knockout luteinized unruptured follicles following PMSG + HCG stimulation (white arrows). (E) LH levels during proestrus at 5 months of age. (F) FSH levels during proestrus at 5 months of age. (G) LH to FSH ratios during proestrus at 5 months of age.

## Notes

*Grant support*: This work was supported in part by the NIH P20 GM121298 (COBRE for Reproductive Health) grant as well as funding from Swim Across America.

### Competing Interest Statement

The authors have declared no competing interest.

### Summary of Updates

The introduction has been shortened and made more relevant to the findings; the methods have additional clarifying details as well as animal numbers (which have been moved from the results section); results have been updated to include new experiments (histology on stimulated ovaries as well as serum FSH and LH measurements); supplemental files have been updated.

