## Supplementary figures and images for "The Requirement of Ubiquitin C-Terminal Hydrolase L1 (UCHL1) in Mouse Ovarian Development and Fertility"

### Supplemental Figure 1

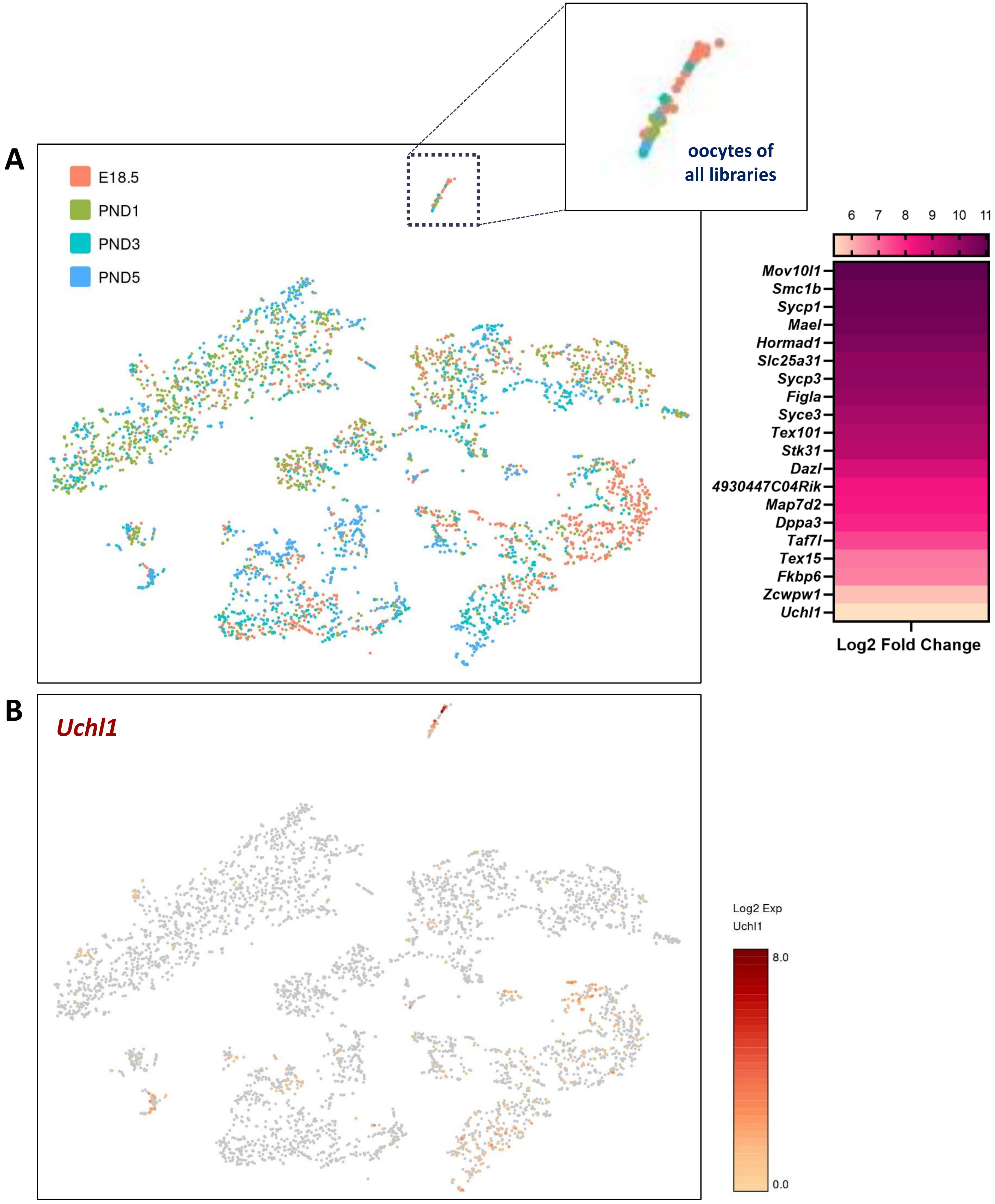

### Supplemental Figure 7

A

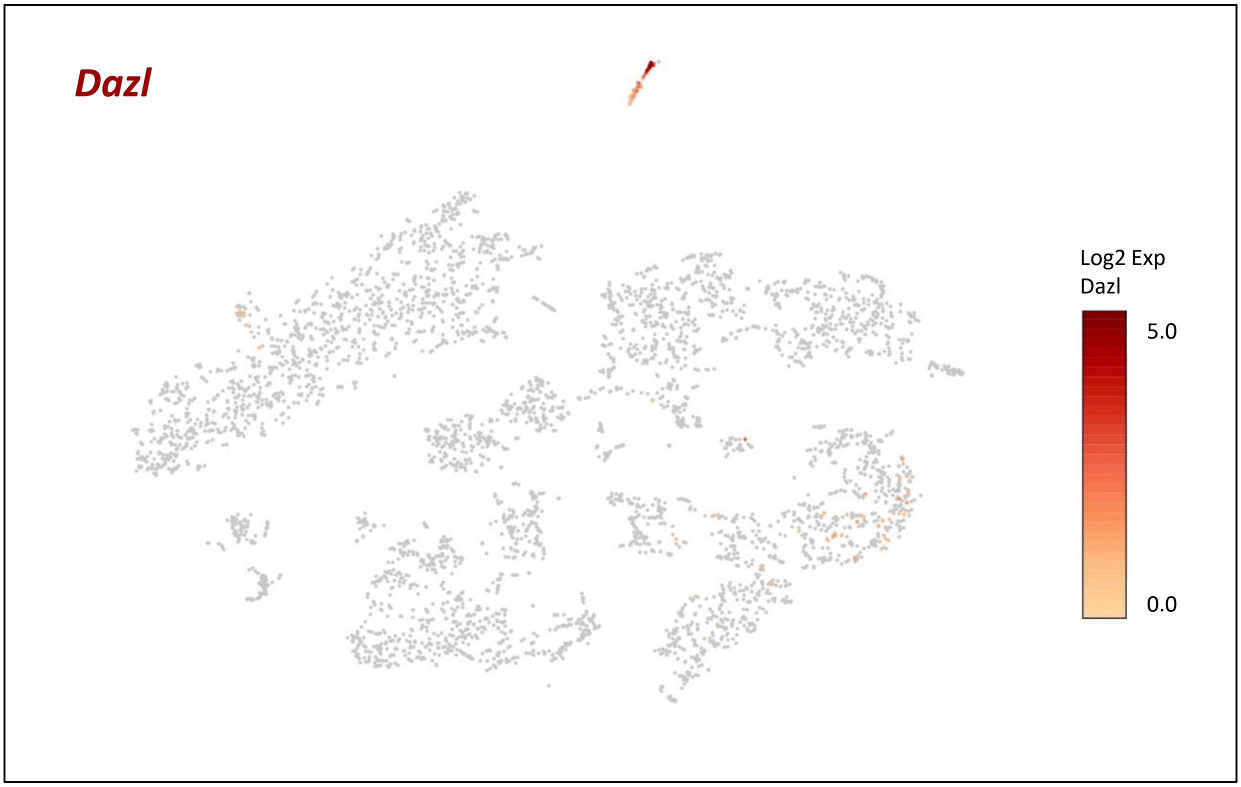


B

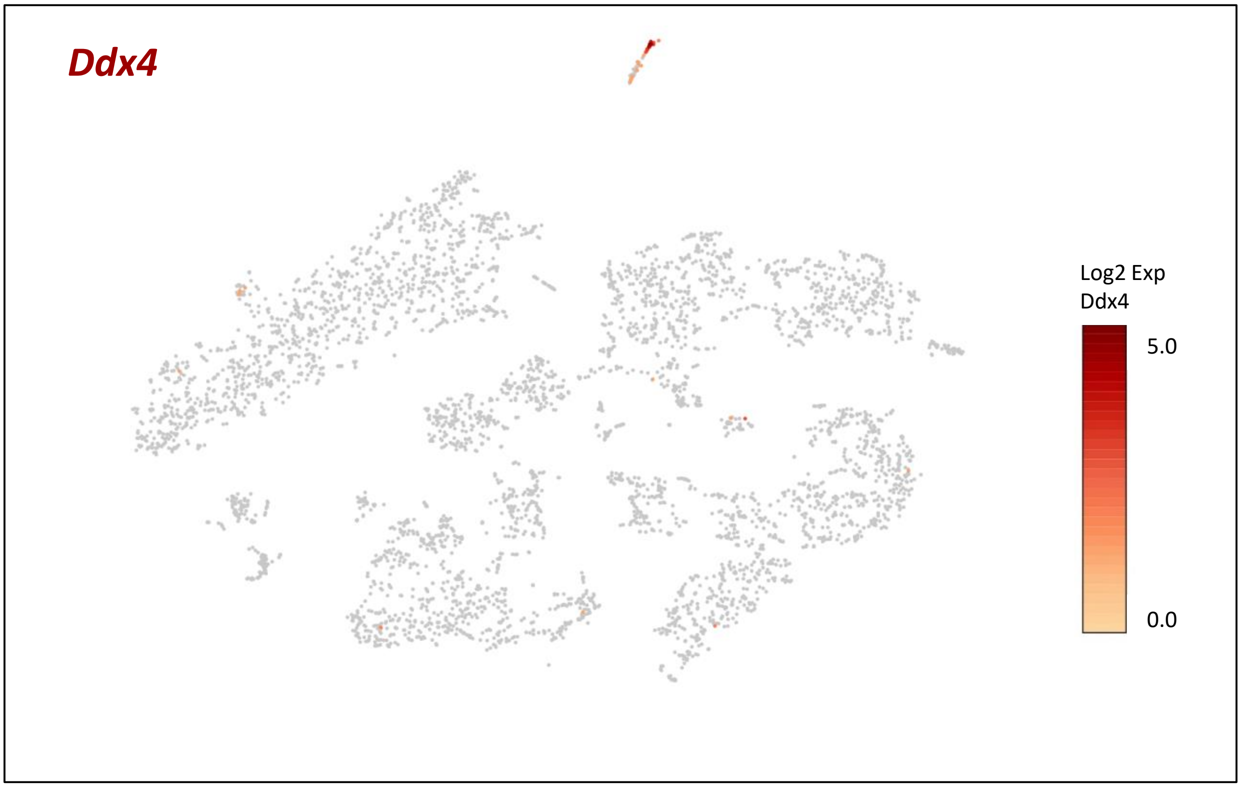


Supp 7

### Supplemental Figure 8

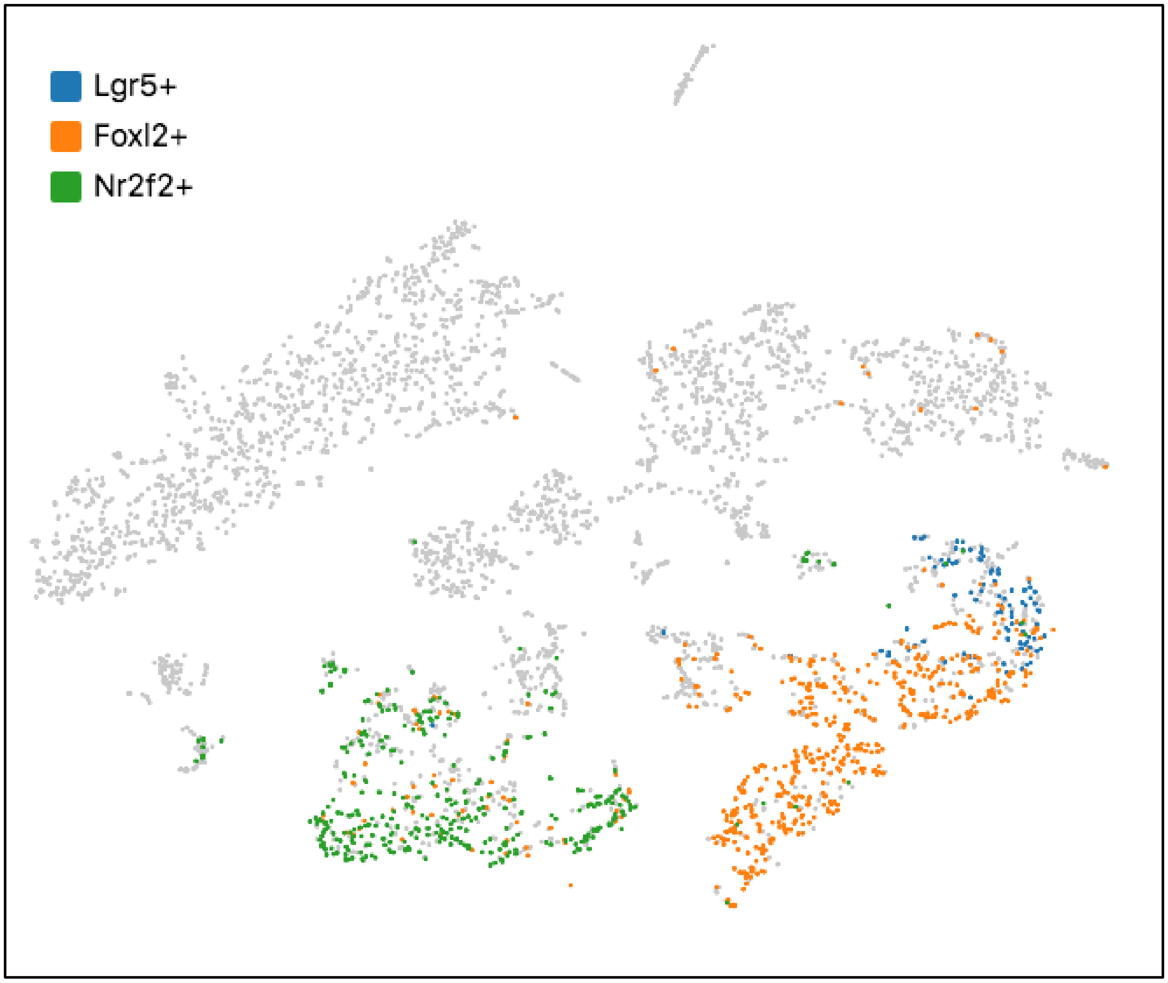


Supp 8

### Supplemental Figure 9

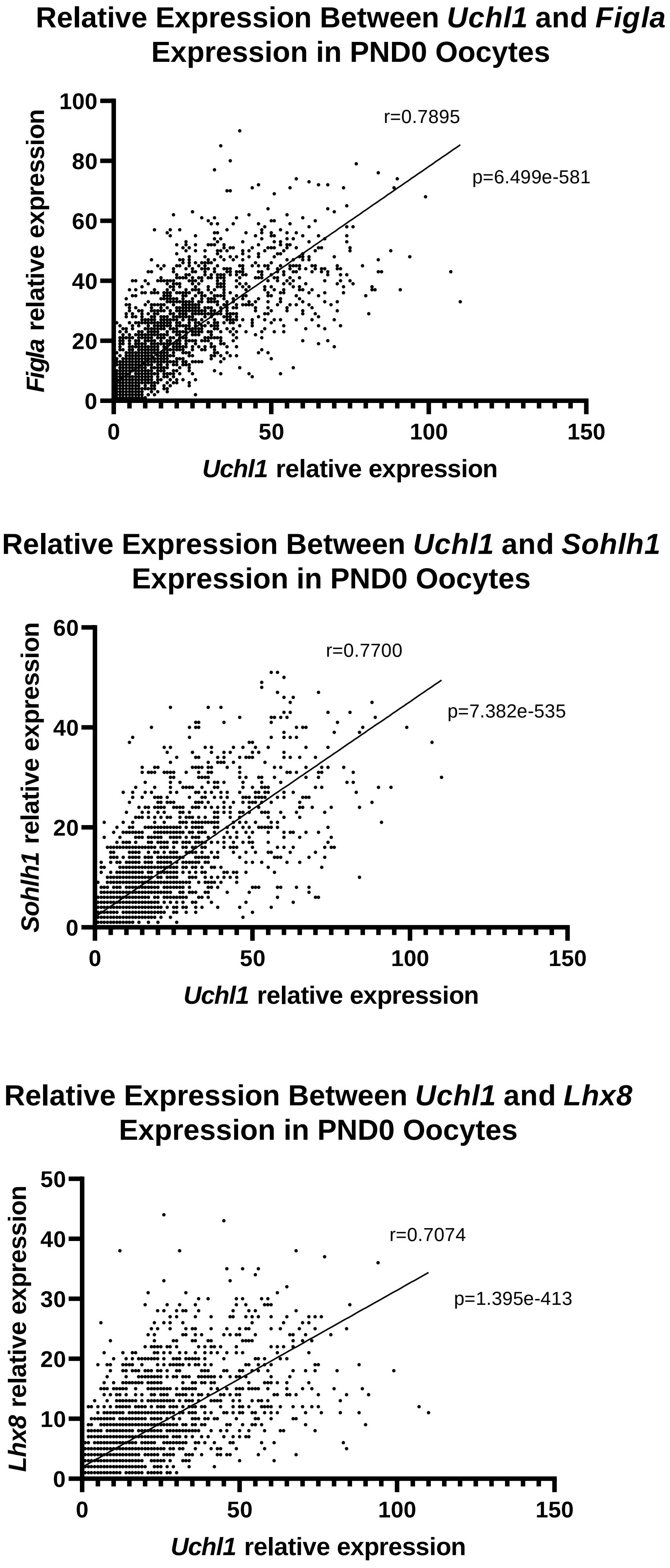

### Supplemental Figure 10

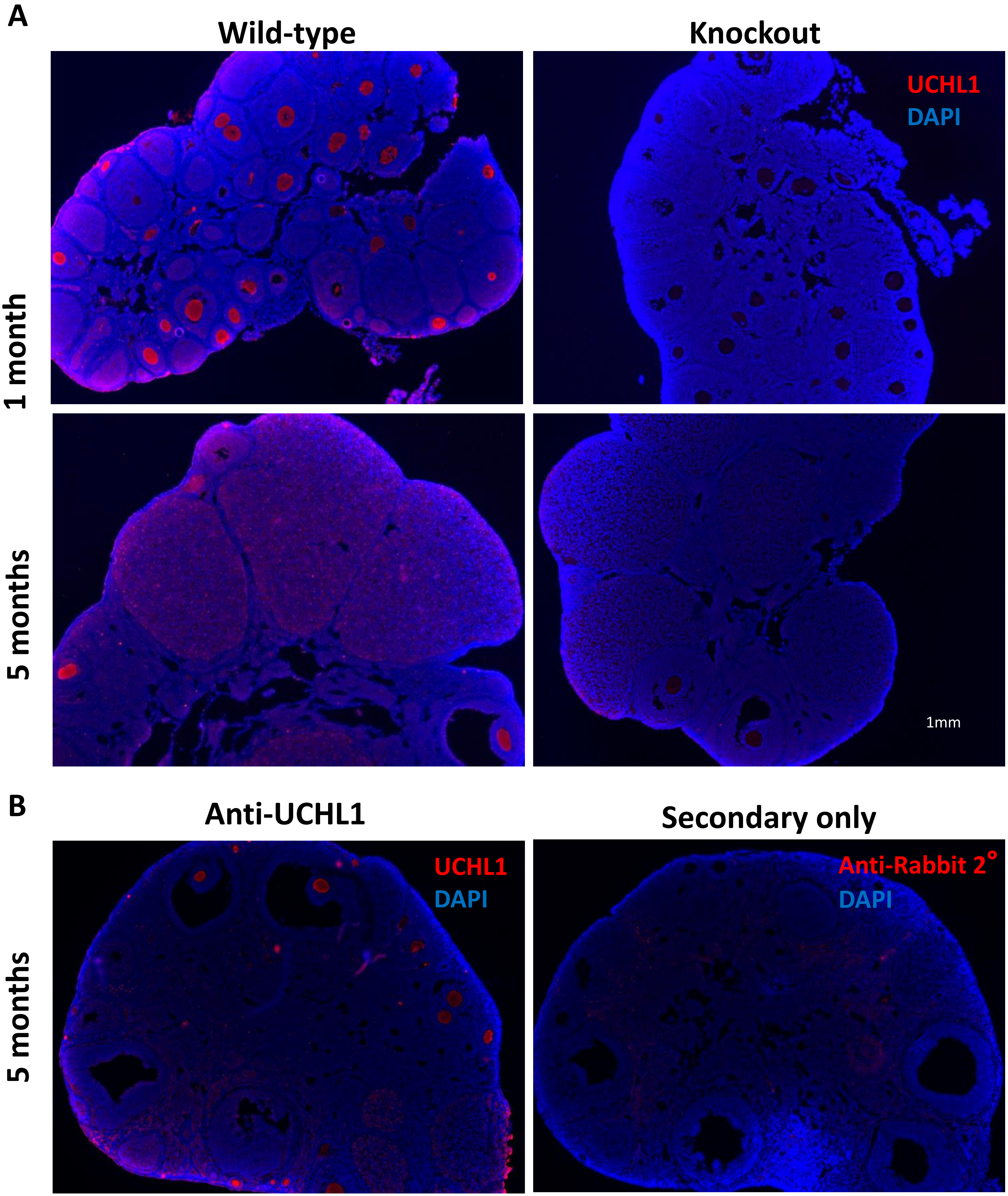

### Supplemental Figure 11

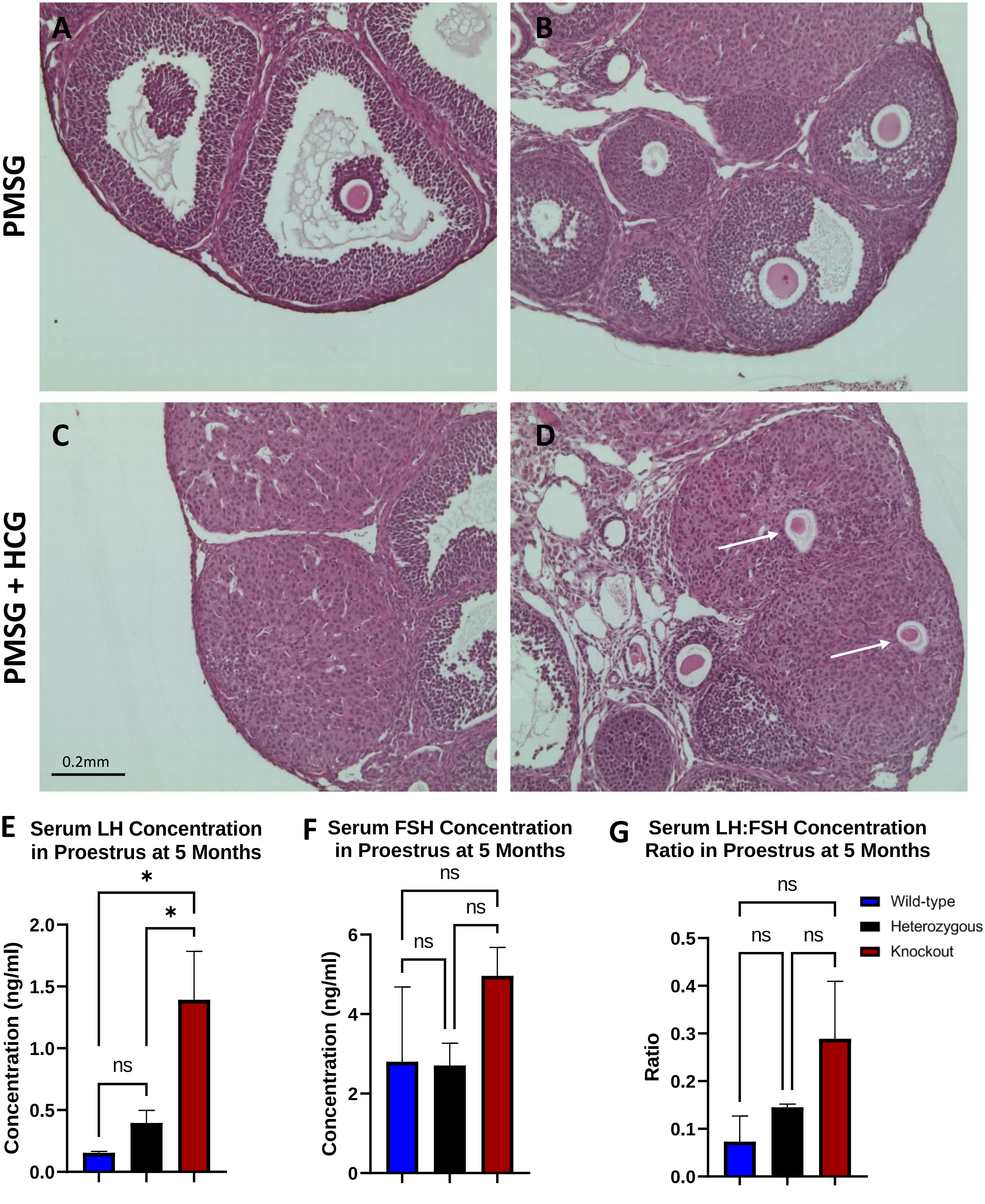
