## Supplemental Figure 2 for "The Requirement of Ubiquitin C-Terminal Hydrolase L1 (UCHL1) in Mouse Ovarian Development and Fertility"

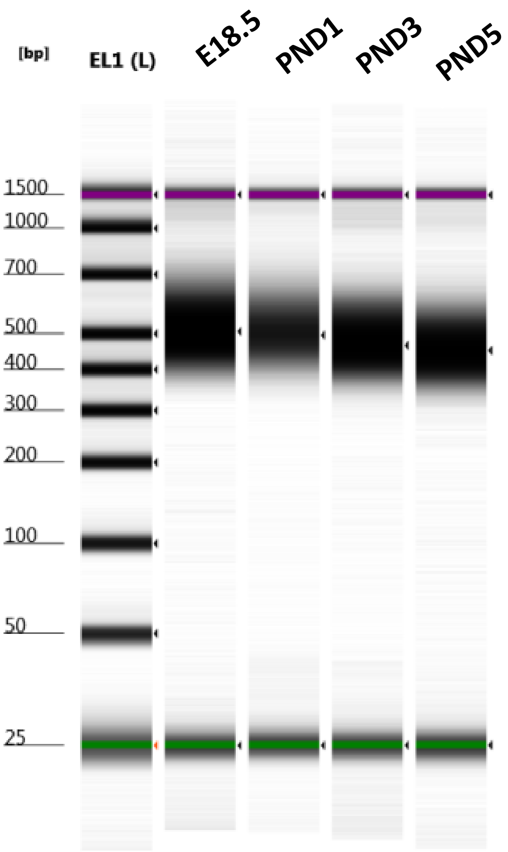


Default image (Contrast 100%)

### Sample Info

| Well | Conc. [pg/µl] | Sample Description | Alert | Observations |
|---|---|---|---|---|
| EL1 | 2350 | Electronic Ladder | | Ladder |
| A1 | 2480 | XCC16833 | | |
| B1 | 1440 | XCC16834 | | |
| C1 | 2380 | XCC16835 | | |
| D1 | 2210 | XCC16836 | | |
