## Supplemental Figure 4 for "The Requirement of Ubiquitin C-Terminal Hydrolase L1 (UCHL1) in Mouse Ovarian Development and Fertility"

**Sample: PND1**

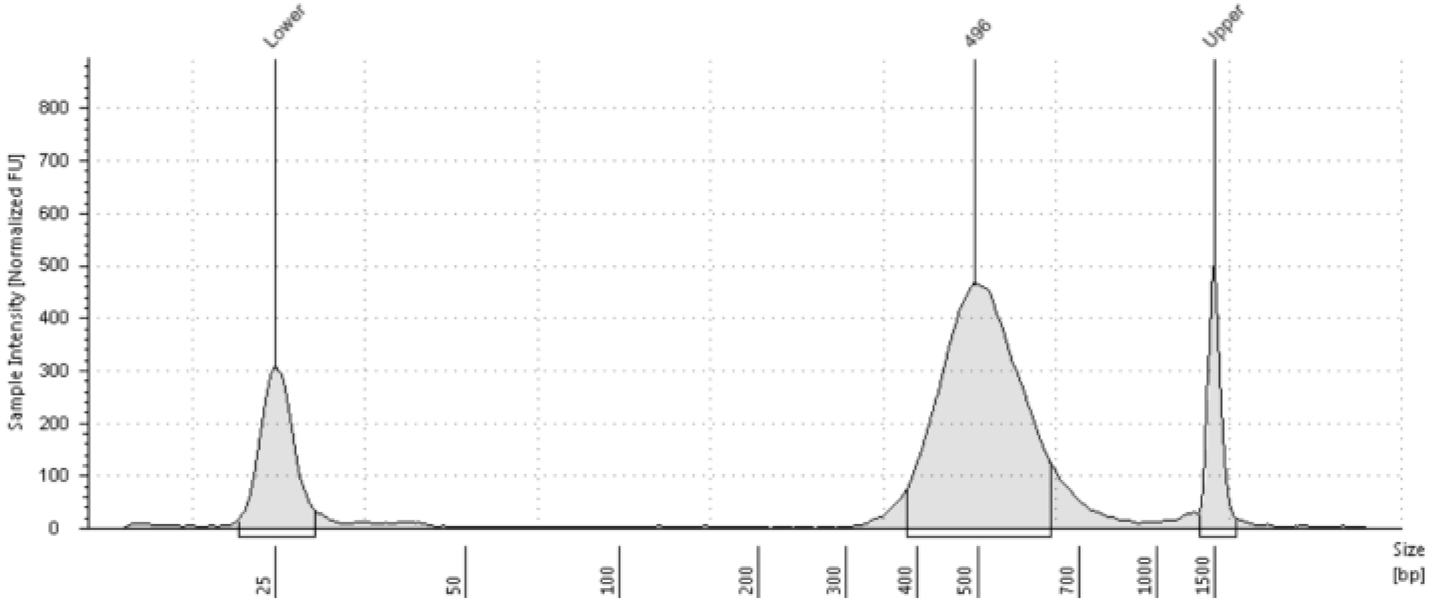


**Sample Table**

| Well | Conc. [pg/µl] | Sample Description | Alert | Observations |
|---|---|---|---|---|
| B1 | 1440 | XCC16834 | | |

**Peak Table**

| Size [bp] | Calibrated Conc. [pg/µl] | Assigned Conc. [pg/µl] | Peak Molarity [pmol/l] | % Integrated Area | Peak Comment | Observations |
|---|---|---|---|---|---|---|
| 25 | 399 | - | 24500 | - | | Lower Marker |
| 496 | 1440 | - | 4480 | 100.00 | | |
| 1500 | 250 | 250 | 256 | - | | Upper Marker |

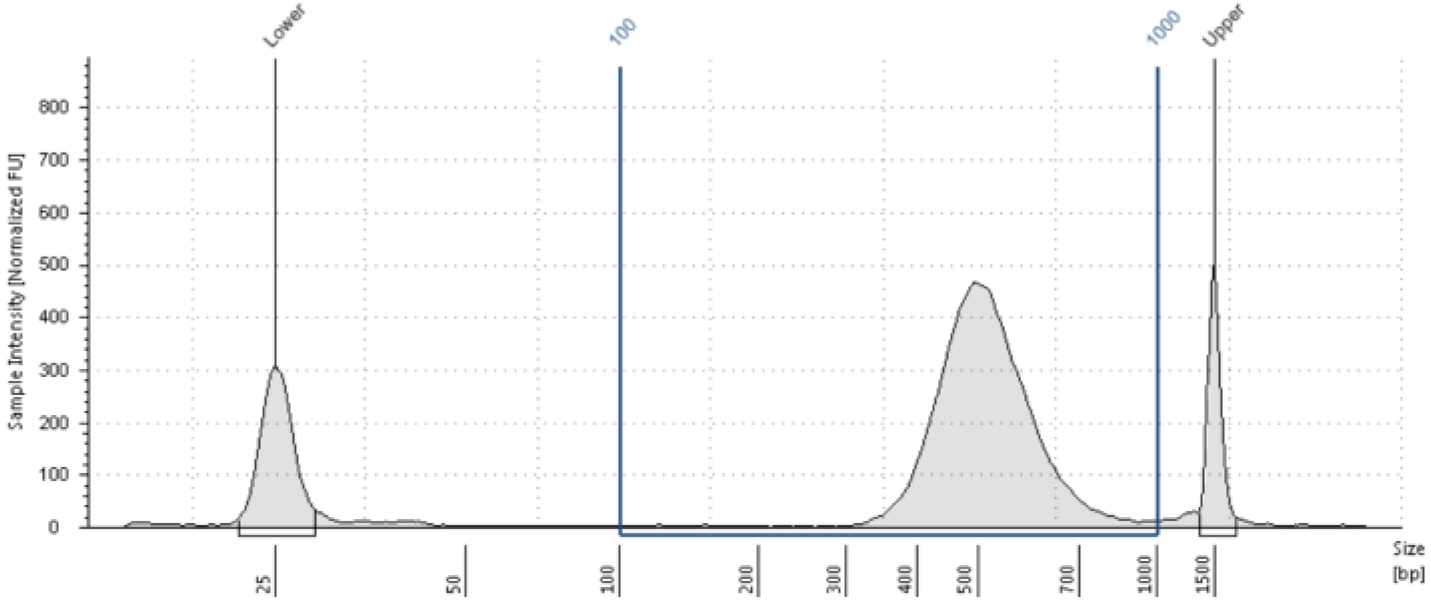


**Region Table**

| From [bp] | To [bp] | Average Size [bp] | Conc. [pg/µl] | Region Molarity [pmol/l] | % of Total | Region Comment | Color |
|---|---|---|---|---|---|---|---|
| 100 | 1000 | 523 | 1620 | 5000 | 94.54 | | |

Supp 4
