## Supplemental Figure 6 for "The Requirement of Ubiquitin C-Terminal Hydrolase L1 (UCHL1) in Mouse Ovarian Development and Fertility"

**Sample: PND5**

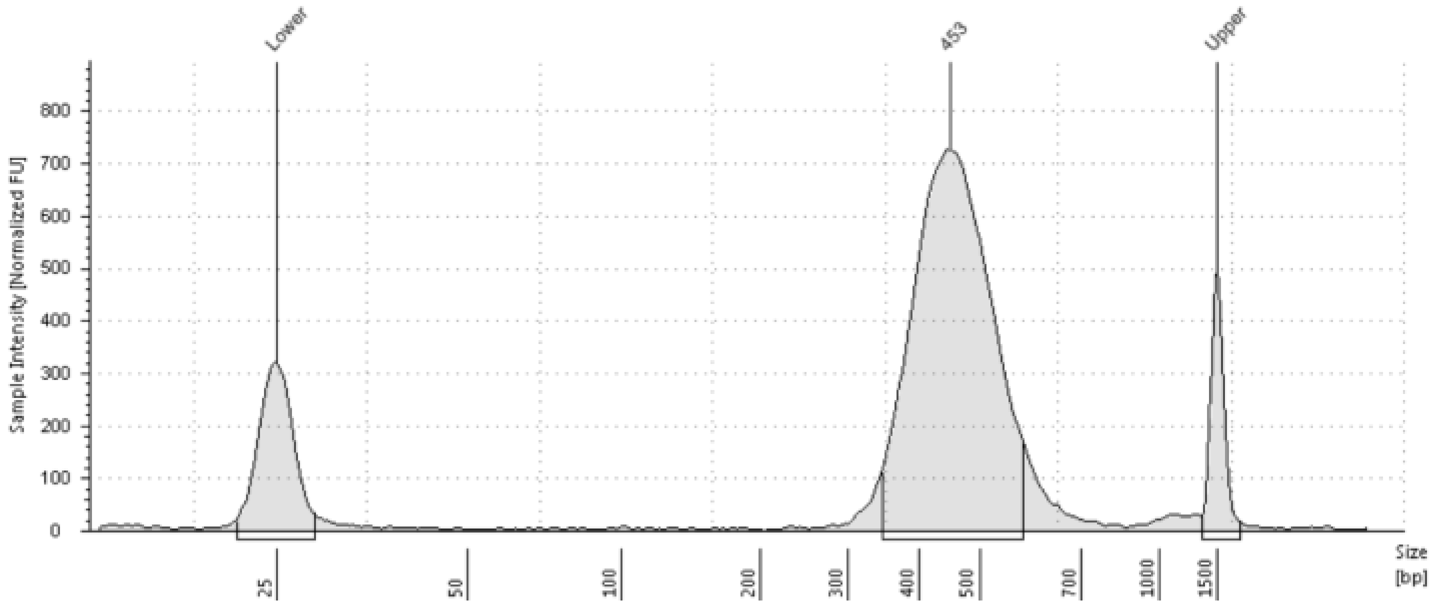


**Sample Table**

| Well | Conc. [pg/µl] | Sample Description | Alert | Observations |
|---|---|---|---|---|
| D1 | 2210 | XCC16836 | | |

**Peak Table**

| Size [bp] | Calibrated Conc. [pg/µl] | Assigned Conc. [pg/µl] | Peak Molarity [pmol/l] | % Integrated Area | Peak Comment | Observations |
|---|---|---|---|---|---|---|
| 25 | 437 | - | 26900 | - | | Lower Marker |
| 453 | 2210 | - | 7510 | 100.00 | | |
| 1500 | 250 | 250 | 256 | - | | Upper Marker |

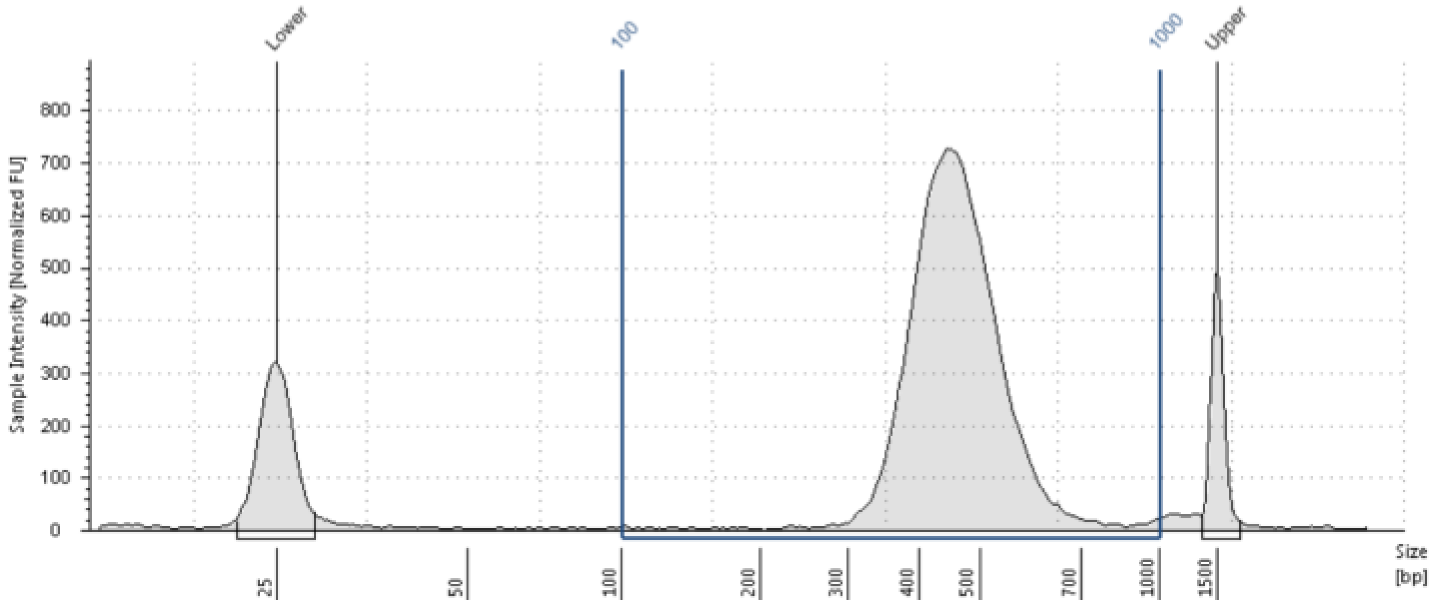


**Region Table**

| From [bp] | To [bp] | Average Size [bp] | Conc. [pg/µl] | Region Molarity [pmol/l] | % of Total | Region Comment | Color |
|---|---|---|---|---|---|---|---|
| 100 | 1000 | 468 | 2450 | 8440 | 95.61 | | ■ |

Supp 6
